# Functional characterization of bat limb regulatory elements

**DOI:** 10.64898/2026.04.07.717074

**Authors:** Aki Ushiki, Guy Kelman, Rory Sheng, Elizabeth Murray, Walter Eckalbar, Yichi Zhang, Mai Nobuhara, Riya Rajani, Katja Friess, Vladyslav Barskyi, Kelly Ngo, Shiori Kinoshita, Stephen A Schlebusch, Mandy Mason, Shijun Zhan, Maggie Liang, Sarah Fong, Md Yasin Haider, Varda Singhal, Tony Schountz, Dorit Hockman, Nicola Illing, Tommy Kaplan, Nadav Ahituv

## Abstract

Bats are the only mammals capable of powered flight and roost head-down. However, the molecular changes shaping bat limbs remain largely unknown. Here, we used comparative functional genomics coupled with mouse-bat sequence swaps to identify key regulatory elements important in bat limb development. We generated and compared bat and mouse forelimb and hindlimb genomic datasets at key wing developmental timepoints, followed by mouse enhancer assays to characterize sequences showing differences between species. We then swapped six mouse enhancer sequences with their corresponding bat sequences, obtaining a variety of bat limb associated phenotypes, including ossification delay, longer digits, thicker skin and symmetrical hindlimb digits. Our work provides a genomic catalog of genes and regulatory elements involved in bat limb development and through extensive characterization in mice shows how changes in regulatory elements lead to small phenotypic changes that together contribute to bat limb development.

## Main

Bats are the only mammal capable of powered flight and roost head-down. To achieve this, they have undergone a variety of morphological innovations, that include not only the wing itself but also unique muscles that power it along with higher metabolic rates, flexible bones, the lack of claws on digits II to V in the forelimb, and craniofacial changes such as having a smaller premaxilla and changes in ear placement to reduce drag^1^. In terms of the wing, during development the bat forelimb autopods retain extended embryonic growth of the metacarpals and proximal and intermediate phalanges of digits II to V^1–4^. In addition, developing bat autopods retain their interdigital tissue which forms part of the wing^1,5^. In contrast, the bat hindlimbs have relatively short, free, symmetrical digits. The hindlimb is diminutive in comparison, rotated by 180°, with short symmetrical clawed digits that allows bats to lock themselves into a roosting position with a tight grip^6,7^.

Examining gene expression differences between bat forelimb and hindlimb at key embryonic developmental stages has found numerous genes to be associated with bat wing development. This was initially carried out in a ‘gene-by-gene’ manner, finding genes such as *Tbx3*, *Brinp3, Meis2*, *HoxD* genes and members of the *Shh-Fgf* signaling pathway to be associated with bat-specific changes in forelimb development^5,8–11^. Later studies used genome-wide assays, such as RNA-seq, on developing bat forelimbs and hindlimbs at key wing developmental stages finding a multitude of gene expression changes between forelimbs and hindlimbs, primarily in known limb developmental associated genes^12,13^. These include multiple ribosomal proteins and known limb patterning signaling pathways, including FGF, WNT, and BMP signaling. Recent work that used single-cell RNA-seq on the bat wing found a specific fibroblast cell population in the bat wing membrane that expresses *MEIS2* and *TBX3* that is thought to be important in tissue retention^14,15^ and a unique bat forelimb mesenchymal progenitor cell population with high *PDGFD* expression that is thought to promote bone proliferation^15^. Combined, these studies suggest that changes in regulatory elements that differentially activate these genes in the forelimb are major drivers in the evolution of the bat wing.

Gene regulatory elements, such as promoters and enhancers, control spatial temporal and levels of gene expression. Several individual enhancers associated with bat wing development have been investigated on a ‘one-by-one’ basis. One such example is the *Prx1* limb enhancer. The replacement of this enhancer with the homologous bat *Prx1* sequence resulted in mouse embryonic day (E) 18.5 embryos having 6% longer forelimbs^16^. Using comparative genomics to annotate bat accelerated regions (BARs), sequences that are conserved in mammals but significantly changed in bats, identified hundreds of candidate regions^13,17,18^. Mouse enhancer assays using the bat sequence of five of these BARs, chosen due to their proximity to known limb genes, found all five to be limb enhancers and three to have differential enhancer activity compared to their homologous mouse sequence^17^. In addition, using chromatin immunoprecipitation followed by sequencing (ChIP-seq) for active [H3K27ac;^19,20^] and repressive [H3K27me3;^21^] marks on autopods dissected from bat forelimbs and hindlimbs at key wing developmental stages (CS15, CS16 and CS17) from the Natal long-fingered bat (*Miniopterus natalensis*), identified thousands of differential active regulatory elements^13^. Combined, these studies have identified thousands of candidate genes and regulatory elements involved in this morphological adaptation, making it difficult to pinpoint the sequences driving this adaptation.

We took a variety of approaches to narrow down the number of candidate genes and regulatory elements specifically involved in bat wing development for further studies including: 1) expression/activity differences from genomic datasets; 2) focusing on biologically relevant genes; 3) BAR selection. We first generated genomic datasets (RNA-seq, H3K27ac & H3K27me3 ChIP-seq) for mouse forelimb and hindlimb for stages of development that match bat embryonic stages of development (CS15-CS17)^22^. In addition, we generated ATAC-seq datasets for developing bat forelimb and hindlimb samples at CS17and embryonic day (E) 13.5 embryos. Utilizing these datasets, we were able to tease out genes and candidate regulatory sequences that specifically changed in the bat forelimbs during these key stages of wing development. We next carried out mouse enhancer assays for eleven candidate bat enhancer sequences, finding several functional limb enhancers that show differential activity compared to their homologous mouse sequences. To characterize their functional importance, we next swapped six sequences in the mouse with the homologous bat sequence, finding a multitude of phenotypes associated with bat limb development, including delayed bone ossification, longer digits, thicker skin and symmetrical hindlimb digits. Combined, our work highlights specific genes and regulatory regions important for bat wing development. In addition, by functionally linking these elements to different phenotypic outcomes through extensive characterization in mice, we show how changes in regulatory elements lead to small phenotypic changes that together contribute to bat limb development in line with recent hypotheses that large evolutionary changes are caused by multiple factors^23^.

## Results

### Transcriptomic comparison of embryonic bat and mouse limbs

Previous work by our labs identified over 7,000 genes to be differentially expressed between forelimb (FL) and hindlimb (HL) in *M. natalensis* during Carnegie Stages 15-17 (CS15-CS17), three key stages of bat wing development^13^. As there are inherent differences in gene expression in FL compared to HL autopods in flightless mammals, we carried out RNA-seq on mouse FL and HL at matching developmental changes, embryonic day (E) 12.0, 13.0 and 13.5 matching CS15, CS16 and CS17 respectively (**Fig. 1a**). We then compared FL and HL differentially expressed (DE) genes in bats and mice, highlighting specific gene expression changes that are unique to bat. We identified a total of 1,715 DE genes between FL and HL. Amongst them, 123 genes (7%) were shared between bats and mice and 1,234 (72%) and 358 (21%) were unique to bat and mouse respectively (**Fig. 1b**). For the bat FL, these included *HoxD* genes (*Hoxd10-13*) and *Tbx3* that showed higher expression in the bat FL compared to mouse (**Fig. 1c-d**). In the bat HL, we observed *HoxC* genes (*Hoxc8-10*) and *Tbx4* to have higher expression compared to mouse (**Fig. 1c-d**). Gene ontology (GO) enrichment analyses using DAVID^24^ for the bat differential genes (N=1,234) found enrichment for terms that are involved in development (**Extended Data Fig. 1a**, **Supplementary Data 1**). Comparison of GeneORGANizer^25^ results for bat only (N=1,234) versus mouse only (N=358) DE genes found an enrichment for bat DE genes known to be expressed in the skeleton (**Extended Data Fig. 1b**), including ones specific to the digits (**Extended Fig. 1c**).

**Fig. 1:**
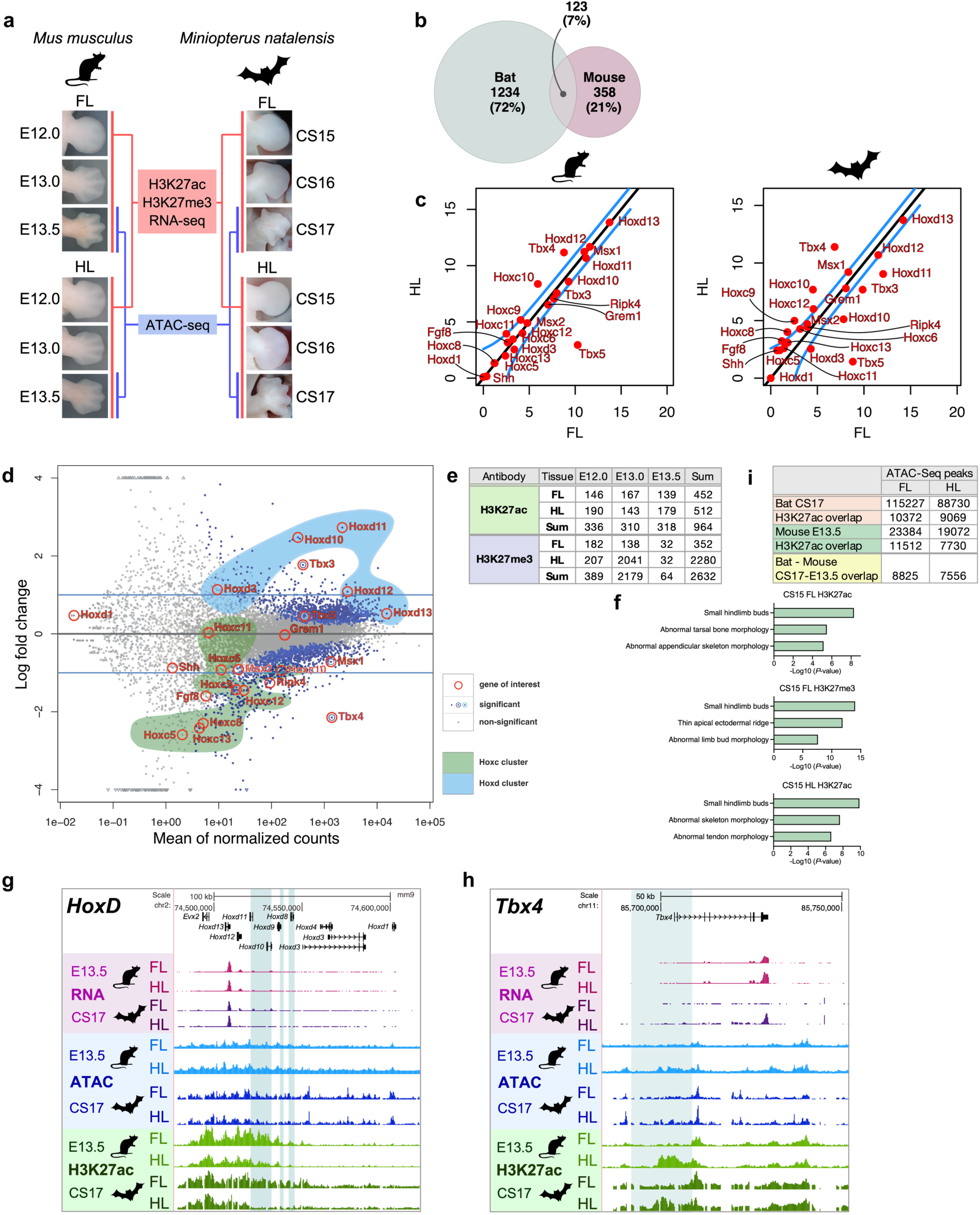
Comparison of RNA-seq and ChIP-seq data between mouse and bat. **a**, Experimental Design. RNA-seq and ChIP-seq (H3K27ac and H3K27me3) were carried out on autopods from *forelimb* (FL) and *hindlimb* (HL) at three developmental stages (E12.0, E13.0, and E13.5 in mouse and the corresponding CS15, CS16 and CS17 in bat). ATAC-seq was conducted at E13.5 and CS17. **b**, Venn diagram showing the number of differentially expressed genes between FL and HL in bats and mice. **c**, Comparison of FL vs. HL differentially expressed genes in mice (left panel) and bats (right panel). The X and Y axes are normalized read counts. The blue lines reference a log2-fold change of 1, i.e. deviation twice away from what would be expected under the assumption that FL and HL expression are identical. **d**, MA plot of differential gene expression between bat and mouse, considering the tissue (FL vs. HL) and stage (E12 through E13.5 and CS15 through C17). The X-axis provides the mean normalized counts, placed against the model’s log2-fold change on the Y-axis. The blue dots mark a statistical significance of this differential gene expression across the species, tissue, and time stages. Similar to panel C, the light blue horizontal lines reference a log2 fold-change of 1. Selected genes are marked by orange circles and their gene symbols are given. Transcripts in the *Hoxc* and *Hoxd* clusters are within the green and blue shades respectively. **e**, FL and HL differentially active (DA) ChIP-seq peak numbers for H3K27ac and H3K27me3 at three stages of development for mouse compared to bat. The stage designation in the headers represent mouse and bat, such that E12.0 is the equivalent bat developmental stage CS15. **f**, GREAT^26^ analysis of H3K27ac and H3K27me3 ChIP-seq DA peaks for CS15 FL and HL autopods, compared to the read count pattern of the corresponding region in the mouse. No limb-related associated terms were found for CS15 HL H3K27me3 and hence not shown. **g**-**h**, DA peaks in the *HoxD* (**g**) and *Tbx4* (**h**) locus with the following tracks: RNA-seq (pink), ATAC-seq (blue), H3K27ac ChIP-seq (green) for E13.5 mouse and CS17 bat. The blue highlighted regions indicate DA peaks. The Y-axis is similar for FL and HL in all tracks. **i**, ATAC-seq peaks in bats (CS17) or mice (E13.5) and their overlap with H3K27ac ChIP-seq and each other.

### Comparison of embryonic bat and mouse limb regulomes

We previously used ChIP-seq to characterize differentially active (H3K27ac) and silenced (H3K27me3) regulatory elements in the three *M. natalensis* developmental stages, CS15, CS16, and CS17^13^. We identified 4,553 and 19,352 differentially enriched regions for H3K27ac and H3K27me3 respectively and 2,475 regions to be differentially enriched between FL and HL for both H3K27ac and H3K27me3^13^. To reduce the number of potential regulatory element candidates involved in wing development, we carried out a similar ChIP-seq for H3K27ac and H3K27me3 in matching mouse FL and HL developmental timepoints (E12.0, E13.0, E13.5) (**Fig. 1a**). To directly compare ChIP-seq peaks between bat and mouse, we aligned our bat peaks to the mouse genome, using only sequences that aligned to a similar location in the mouse genome (see Methods). We then characterized differentially active (DA) peaks between bat and mouse in FL and HL at each developmental time point for either chromatin mark (H3K27ac or H3K27me3), identifying hundreds of DA peaks for each condition (**Fig. 1e**).

We next investigated differentially active (DA) peaks between FL and HL and bat and mouse. Using the Genomic Regions Enrichment of Annotations Tool (GREAT^26^), we analyzed each time point (CS15, CS16, CS17), tissue (FL or HL), and mark (H3K27ac or H3K27me3) individually for enriched phenotype terms. We found enrichment for terms such as small hindlimb buds, abnormal tarsal bone morphology, abnormal appendicular skeleton morphology, thin apical ectodermal ridge, abnormal limb bud morphology, abnormal skeleton morphology, and abnormal tendon morphology (**Fig. 1f**, **Extended Data Fig. 1d**).

To refine the length of candidate sequences, as histone ChIP-seq peaks are quite lengthy, we also carried out ATAC-seq, which provides narrower defined peaks. For bat, this was done on CS17 FL and HL as we only had available *M. natalensis* embryos for this time point and for mouse at E13.5 to match that developmental time point. We identified thousands of peaks in each condition and annotated those that overlap H3K27ac ChIP-seq at a similar time point and that overlap between mouse and bat (**Fig. 1g**). We used our ATAC-seq data as an additional layer to choose candidates for our subsequent mouse assays. We next analyzed individual bat FL DA peaks. We first searched for clusters of DA peaks near FL and HL DE genes. We found gene regulatory DA peak clusters near the *HoxD* genes (*Hoxd10-12*)(**Fig. 1h**) and *Tbx3* for FL and *HoxC* genes and *Tbx4* for HL (**Fig. 1i**), suggesting that these loci are important regions for bat wing and hindlimb development respectively. *Tbx3* and *HoxC* loci were analyzed further in the next results section.

### Functional validation of candidate regions from computational analysis

We next selected three top ranking DA sequences between bat and mouse FL from our genomic assays to characterize for mouse enhancer activity. We chose the following three regions: 1) A previously characterized mouse E11.5 limb enhancer (mm1417, VISTA Enhancer Browser^27^) in the homeobox C cluster (*HoxC*) which has high H3K27ac and ATAC-seq peaks in the bat HL (**Fig. 2a**); 2) A sequence downstream to T-box transcription factor 5 (*Tbx5*) which showed strong H3K27ac and ATAC-seq peaks in the bat FL; 3) A previously characterized mouse E11.5 limb enhancer (hs483, VISTA Enhancer Browser^27^) in the *Tbx3* locus that showed strong H3K27ac activity in the bat FL.

**Fig 2:**
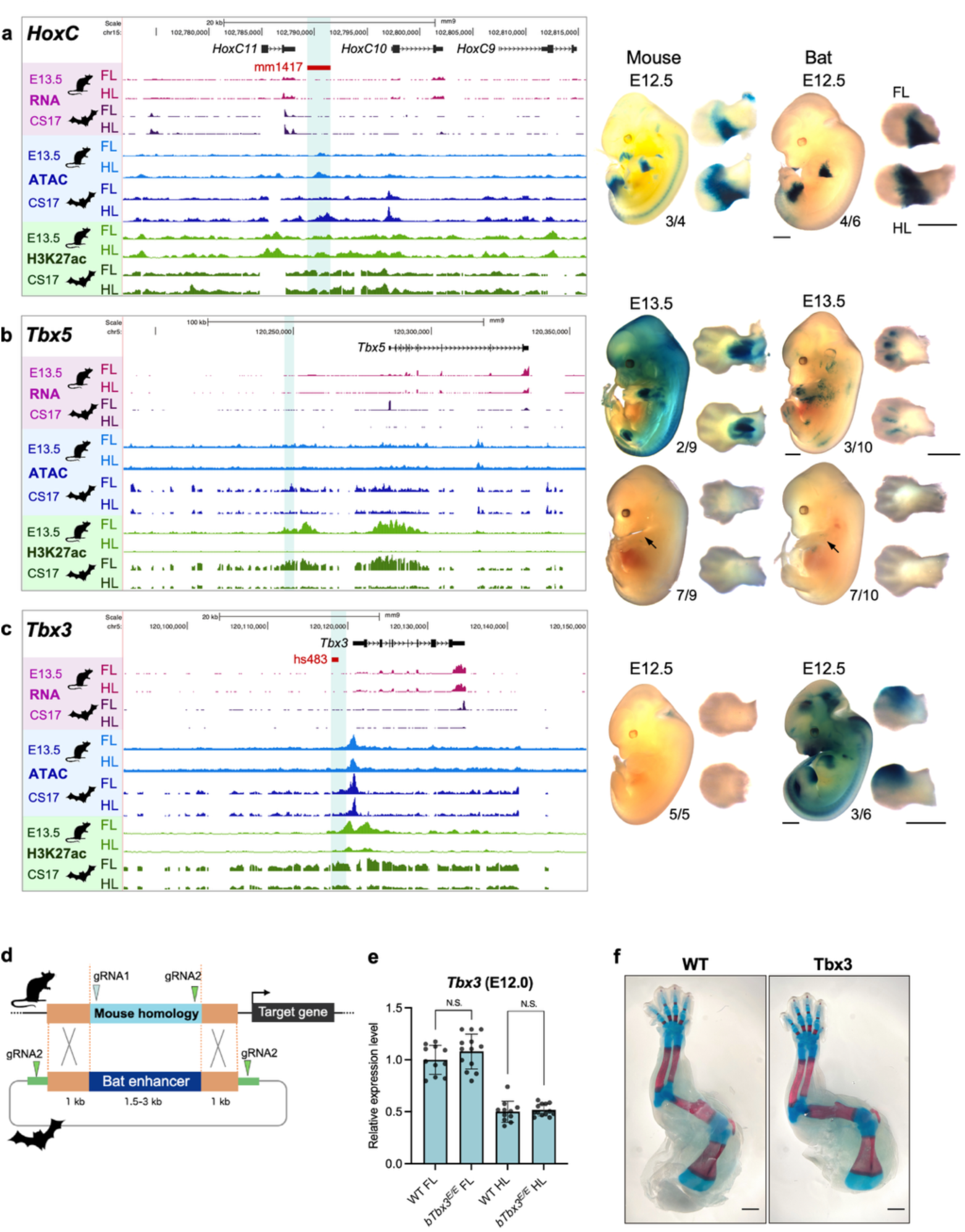
Functional characterization of the bat *HoxC*, *Tbx5* and *Tbx3* enhancers. **a-c,** Genomic landscape and lacZ enhancer transgenic assay. **Left:** RNA-seq (pink), ATAC-seq (blue), and H3K27ac ChIP-seq (green) data from bat or mouse FL and HL at E13.5 (mouse) or CS17 (bat). **Right**: LacZ enhancer transgenic assay result. The number below the representative embryo picture indicates the number of embryos showing similar enhancer activity. Bars below denote 1mm. The *HoxC* locus (**a**) shows mm1417 (red bar in genome snapshot), a validated limb enhancer from the VISTA enhancer browser ^30^. For *Tbx5* (**b**) two distinct staining patterns were observed, with the one above showing strong limb staining and below the black arrows point to the weaker limb staining pattern. For *Tbx3* (**c**) the genome browser shows hs483 (red bar in genome snapshot), a validated limb enhancer from the VISTA enhancer browser ^30^. **d**, The strategy used for bat enhancer replacement. Two gRNAs were designed to target the mouse homologous region (orange) and the donor plasmid is targeted by one of these gRNA (green). **e**, Gene expression levels of *Tbx3* in wild-type (WT) and *bTbx3^E/E^* homozygous knockin mice in FL and HL. Each value represents the ratio of *Tbx3* gene expression to that of *b-Actin* (*n* = 11–13 biological replicates), and values are mean ± standard deviation. The expression value of wild type (WT) FL was arbitrarily set at 1.0. Each dot represents one embryo, and statistical differences were determined using unpaired t test (N.S., not significant). **f**, Alcian blue and alizarin red staining for wild-type (WT) and *bTbx3^E/E^* homozygous knockin mice. Bars below denote 1mm.

All three *M. natalensis* bat sequences were cloned into a mouse enhancer assay vector in front of the *Hsp68* minimal promoter and LacZ reporter gene. In addition, we also cloned the corresponding mouse sequences for the *HoxC*, *Tbx5* and *Tbx3* regions. We observed differences in limb enhancer activity for all three bat sequences, compared to their corresponding mouse sequences at the specified embryonic stage (E12.5 to E13.5) (**Fig. 2a-c**, **Extended Data Fig. 2a-c**). For *HoxC*, we observed limb enhancer activity at E12.5 in the proximal posterior side in both FL and HL for both mouse and bat sequences (**Fig. 2a**). For *Tbx5*, in some embryos we found that the bat sequence had strong enhancer activity in the developing digits while the mouse sequence showed limb enhancer activity in the base of the zeugopod (**Fig. 2b top**); however most embryos had weak enhancer activity in the zeugopod for both bat and mouse sequences (**Fig. 2b bottom**). For *Tbx3*, we observed weak enhancer activity for the mouse sequence while the bat sequence showed strong enhancer activity in the anterior side of the autopod (**Fig. 2c**).

Due to the strong enhancer activity differences between the bat and mouse *Tbx3* sequences and the known role of this gene in early limb development^28^, we swapped the mouse sequence with the bat sequence using a modified protocol for the Precise Integration into Target Chromosome (PITCh) system^29^ that uses longer homology arms (1 kb) and two gRNAs targeting the mouse homologous sequence and the bat enhancer donor plasmid (**Fig. 2d**; see Methods for more detail). We next checked the endogenous *Tbx3* gene expression level at E12.0 limb buds, as this time point is similar to the one we observed enhancer activity differences between the bat and mouse sequence. We observed no significant *Tbx3* expression differences between wild type (WT) and *bat-Tbx3 enhancer* homozygous replacement mice(*bTbx3^E/E^*) (**Fig. 2e**). We also checked for skeletal differences at E18.5 using alcian blue and alizarin red staining, finding no apparent differences (**Fig. 2f**). In summary, despite the strong DA peak differences and transgenic enhancer differences between bat and mouse for this sequence, we did not observe any apparent *Tbx3* expression changes at E12.0 or skeletal differences at E18.5.

### Functional validation of candidate regions from biologically relevant genes

We selected additional enhancer candidates for mouse enhancer assays, based on their known involvement in wing development and/or webbing. These included a sequence near the receptor interacting serine/threonine kinase 4 (*Ripk4*) gene that had strong bat ATAC-seq and H3K27ac peaks in the FL and HL compared to mouse (**Fig. 3a**). Mutations in *RIPK4* cause Bartsocas-Papas type 1 popliteal pterygium syndrome (BPS1; OMIM #263650), that is characterized by cleft lip and/or palate, webbing of the skin in the knees, elbows and neck, syndactyly and genital and digit abnormalities ^31^. In addition, *Ripk4* homozygous mouse knockouts die at birth due to epidermal differentiation abnormalities, including reduced skin folds, fusion of the tail and hindlimb to the body and fusion of the oral cavity, esophagus and external orifices and a more thick epidermis^32^. *Ripk4* was also found to be differentially expressed in the developing bat plagiopatagium (membrane connecting the bat FL and HL) between an insectivorous bat and omnivorous bat suggesting that it is important in shaping wing structure^33^. We also selected a sequence near the gremlin (*Grem1*) gene, which is known to play an important role in limb development^34,35^ and interdigital membrane retention in several mammals including bats^5^. *Grem1* showed high expression levels in the bat FL compared to HL and the chosen sequence showed a FL specific ATAC-seq peak (**Fig. 3b**). This sequence did not overlap any of the previously characterized *Grem1* limb enhancers^36^.

**Fig 3:**
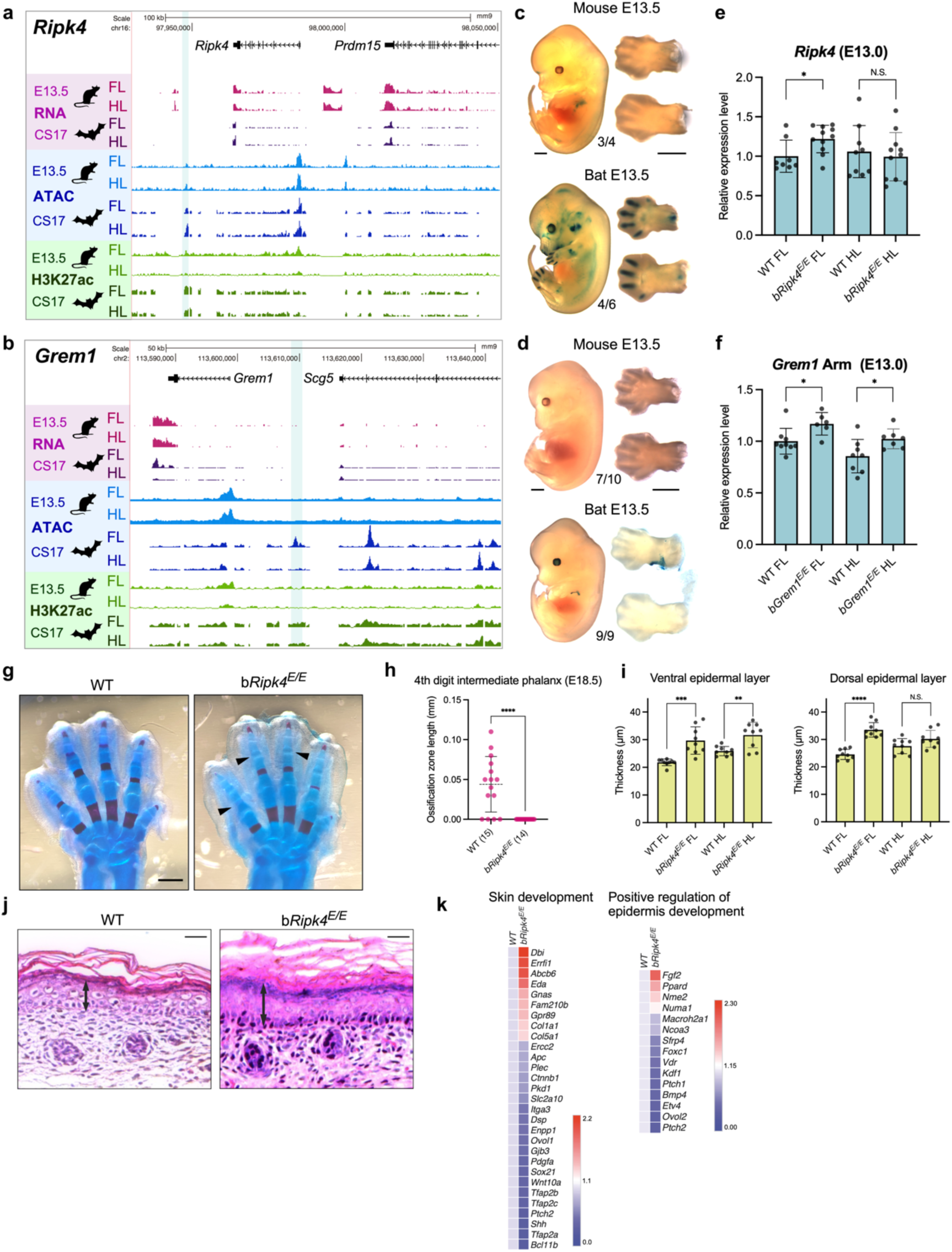
Mouse characterization of bat *Ripk4* and *Grem1* enhancers. **a-f,** Genomic landscape and lacZ enhancer transgenic assay. (**a-b**) RNA-seq (pink), ATAC-seq (blue), and H3K27ac ChIP-seq (green) data from bat or mouse FL and HL at E13.5 (mouse) or CS17 (bat). (**c-d**) LacZ enhancer transgenic assay result. The number below the representative embryo picture indicates the number of embryos showing similar enhancer activity. Bars below denote 1mm. (**e-f**) Gene expression levels of *Ripk4* and *Grem1* in wild-type (WT) and bat enhancer homozygous knockin mice in FL and HL as measured by qRT-PCR. Each value represents the ratio of *Ripk4* and *Grem1* gene expression to that of *b-Actin* (*n* = 6-10 biological replicates), and values are mean ± standard deviation. The expression value of wild type (WT) FL was arbitrarily set at 1.0. Each dot represents one embryo. **g,** Alcian blue and alizarin red staining for wild-type (WT) and *bRipk4^E/E^* homozygous replacement mice. Bars below denote 0.5mm. **h,** Ossification zone length of 4th digit intermediate phalanx at E18.5. Each dot represents one embryo. Biological replicate numbers are shown beneath the graph. **i**, Epidermal skin layer thickness in the FL and HL, measured in both ventral and dorsal regions at P0. Each dot represents one mouse. **j,** H&E staining of FL ventral side skin. Black arrow indicates the epidermis layer. Bars above denote 50μm. **k,** Heat map showing gene expression differences between WT and *bRipk4^E/E^* homozygous replacement mice at P0, based on RNA-seq. Differentially expressed genes associated with the Gene Ontology terms ‘skin development’ or ‘positive regulation of epidermis development’ are shown. All statistical differences were determined using unpaired t test (\**p* < 0.05, \*\**p* < 0.01, \*\*\**p* < 0.005, \*\*\*\**p* < 0.001, N.S., not significant).

We cloned both sequences into a mouse enhancer assay vector and analyzed their LacZ expression at E13.5, as this was the time point that showed strong DA in the ATAC-seq data. We observed strong reproducible expression in the digits and arm for the bat *Ripk4* associated sequence while the mouse sequence only provided dim arm expression (**Fig. 3c**, **Extended Data Fig. 3a**). For the bat *Grem1* associated sequence, we observed enhancer activity in the limb proximal ventral region, while the mouse sequence was negative for enhancer activity (**Fig. 3d**, **Extended Data Fig. 3b**).

Due to the inherent bat and mouse enhancer expression differences, we next swapped both sequences in the mouse using the modified PITCh protocol^29^. We measured mRNA expression for either gene at E13.0, finding a significant increase in *Ripk4* expression in the FL of the *bRipk4^E/E^* mice in the whole limb bud (**Fig. 3e**). We analyzed the *Grem1* expression in autopod (hand) and stylopod/zeugopod (arm) separately.

*Grem1* was significantly upregulated in stylopod/zeugopod; however there was no significant difference in the autopod between the *Grem1* homozygous knockin (*bGrem1^E/E^*) and wild-type mice (**Fig. 3f**, **Extended Data Fig. 3c**). We next characterized bone development at E18.5 for both mouse lines using alcian blue and alizarin red staining, observing no observable differences between *bGrem1^E/E^* and wild-type mice (**Extended Data Fig. 3d**). In *bRipk4^E/E^*mice, we observed a complete absence of ossification in the intermediate phalanges compared to wild-type mice (**Fig. 3g-h**). In bats, delayed ossification in the FL was observed as a means to allow longer digit length^2–4^.

As both *Ripk4* and *Grem1* have important roles in skin development, we next analyzed the ventral and dorsal skin region of the limb of homozygous knockin and wild type mice at P0 using hematoxylin and eosin (H&E) staining. We observed no significant differences in skin epidermis thickness for *bGrem1^E/E^* mice compared to wild-type mice (**Extended Data Fig. 3e**). In contrast, *bRipk4^E/E^* mice had a significantly thicker epidermal skin layer in the FL and HL ventral region and in the FL dorsal region (**Fig. 3i-j**).

To further characterize the molecular mechanisms driving this epidermal skin layer length difference, we conducted RNA-seq on *bRipk4^E/E^*and WT P0 FL ventral skin samples (**Fig. 3k**). Gene Ontology (GO)^37,38^ term enrichment for DE genes between *bRipk4^E/E^*and WT mice found significant enrichment for “skin development” and “positive regulation of epidermis development”. One of the top DE genes was the Fos proto-oncogene (*Fos*) that showed 6.7 fold higher expression in *bRipk4^E/E^*mice (**Extended Data Fig. 4**). Inducible expression of *c-fos* in the epidermis of adult mice is known to promote inflammation-mediated epidermal hyperplasia^39^. Other DE genes included the ERBB receptor feedback inhibitor 1 (*Errfi1*, also known as *Mig6i)*, a protein that regulates the epidermal growth factor receptor (EGFR) pathway and is involved in skin development^40^ and the Transcription Factor AP-2 Alpha (*Tfap2a*, also known as *AP-2α*) whose loss leads to altered epidermal morphology^41^. We also found the fibroblast growth factor 2 (*Fgf2*) which binds to the fibroblast growth factor receptor 2 (Fgfr2) in keratinocytes to trigger a signaling cascade that stimulates cell division, promoting epidermal growth and thickness^42^ to be DE. Combined, our RNA-seq results identified numerous DE genes which could contribute to a thickened epidermis layer in *bRipk4^E/E^*.

### Functional validation of bat accelerated regions

We next characterized BARs near genes involved in limb development. These included: 1) BAR116, a sequence in the *Hoxd* locus (**Extended Data Fig. 5a**) that we previously showed to have strong FL expression in the entire autopod at E12.5 while the mouse sequence was negative for enhancer activity^17^. It is also worth noting, that the homologous BAR116 chicken sequence showed increased FL expression compared to the HL in mouse enhancer assays at E12.5^43^. 2) The sonic hedgehog (*Shh*) limb enhancer (BAR61), also called the zone of polarizing activity (ZPA) regulatory sequence (ZRS) or the mammal-fish-conserved-sequence 1 (MFCS1)^44^, did not show significant expression differences between the *M.natalensis* bat and mouse sequences in a transgenic mouse enhancer assay at E12.5^17^ but showed extended activity at the anterior edge of the limb for the *Pteropus Vampyrus* (megabat) sequence at E11.5 compared to mouse^45^. The ZRS is associated with a variety of limb malformations including polydactyly, triphalangeal thumb and syndactyly in humans^46^, limb loss in snakes^45^ and limb malformations in chickens^47^. 3) Two adjacent BARs near the msh homeobox 2 (*Msx2*) gene^18^ that along with *Msx1* plays an important role in mesenchyme interdigit cell death during limb development^48,49^ and both genes were found to have significantly lower expression in the embryonic bat FL compared to the HL^13^.

As mouse enhancer assays for the bat and mouse sequence were previously carried out for BAR116 and ZRS (BAR61), we swapped these two sequences. For BAR116, we did not observe any gene expression differences via qRT-PCR in FL for *Hoxd9*, *10*, *11*, *12*, and*13* at E12.5, the time point where the bat and mouse showed differential enhancer activity^17^, between the homozygous knockin and wild-type mice, but observed some differences in the HL for *Hoxd9*, *Hoxd10*, and *Hoxd12* (**Extended Data Fig. 5b**). However, we did not observe any apparent skeletal differences in the FL and HL at P1 using alcian blue and alizarin red staining (**Extended Data Fig. 5c**). In contrast, for *bZRS^E/E^*, we observed polydactyly in the HL in heterozygous mice that became more symmetrical and 100% penetrant in homozygous mice, similar to a bat HL (**Fig. 4a**). We also observed a shortened tibia (**Fig. 4b**). In the FL, we observed lower levels of mild polydactyly (55% of the mice) (**Fig. 4b**). qRT-PCR on the E11.5 limb showed *Shh* to be significantly upregulated only in the HL (**Fig. 4e**). Whole-mount *in situ* hybridization (WISH) for *Shh* and its receptor patched 1 (*Ptch1*) found both to show ectopic expression in the anterior HL (**Fig. 4c-d**). We also analyzed previous *Shh* WISH on bat *M. natalensis* embryos^8^ at CS14E (equivalent to E11.0 in mice) observing similar ectopic expression in the anterior part of the HL in two different embryos (**Fig. 4f**, **Extended Data Fig. 5d**).

**Fig 4:**
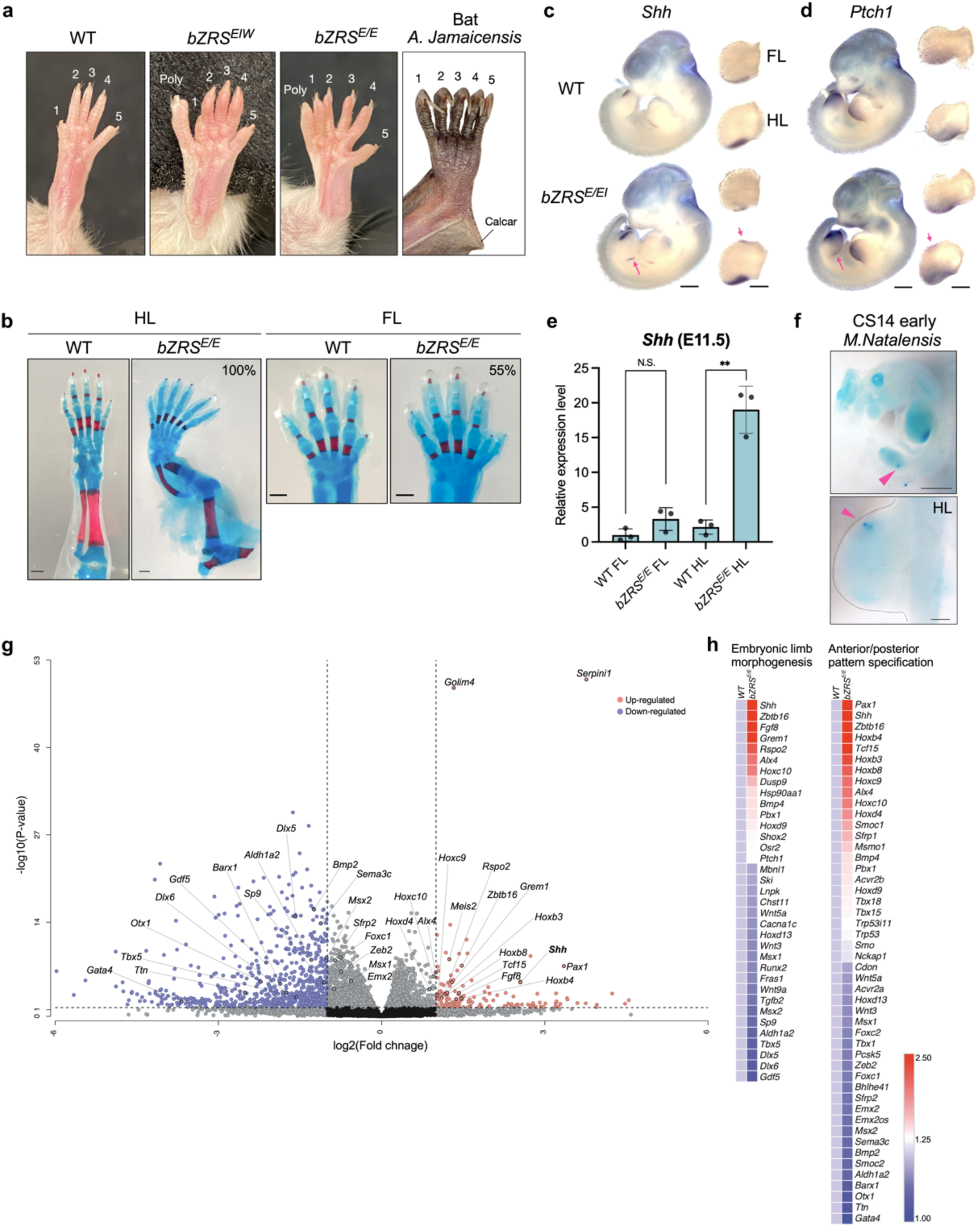
Mouse characterization of bat ZRS. **a,** Hindlimb phenotype in *bZRS^E/E^*. Hindlimb morphology in WT, *bZRS^E/W^*, *bZRS^E/E^*, and *Artibeus jamaicensis*. Polydactylous digits are indicated as poly. **b,** Alcian blue and alizarin red staining for wild-type (WT) and *bZRS^E/E^* homozygous replacement mice. Bars below denote 0.5 mm. Percentage noted on the *bZRS^E/E^* panel is the penetrance of this phenotype. **c-d,** Whole-mount *in situ* hybridization for *Shh* (**c**) and *Ptch1* (**d**) of E11.5 mouse embryos. Black bar represents 1 mm (whole embryo) and 0.5 mm (limb bud). **e,** Gene expression levels of *Shh* in wild-type (WT) and bat enhancer homozygous replacement mice in FL and HL as measured by qRT-PCR. Each value represents the ratio of *Shh* gene expression to that of *b-Actin* (*n* = 3 biological replicates), and values are mean ± standard deviation. The expression value of wild type (WT) FL as arbitrarily set at 1.0. Each dot represents one embryo, and statistical differences were determined using unpaired t test (\*\**p* < 0.01, N.S., not significant). **f,** Whole-mount *in situ* hybridization for *Shh* of CS14 early *Miniopterus natalensis* embryo. Black bar represents 1 mm (whole embryo) and 0.2 mm (limb bud). Pink arrowhead indicates the *Shh* gene expression in HL. **g,** Volcano plots showing the global transcriptional changes for the indicated groups. Each circle represents one gene. The log2 fold change in the indicated genotype is represented on the x axis. The y axis shows the p value. A p value of 0.05 and a fold change of 2 are indicated by lines. **h,** Heat map showing gene expression differences between WT and *bZRS^E/E^* homozygous knock-in mice at E11.5, based on RNA-seq. Differentially expressed genes associated with the Gene Ontology terms ‘Embryonic limb morphogenesis’ or ‘Anterior/posterior pattern specification’ are shown.

We next carried out RNA-seq on both WT and *bZRS^E/E^* E11.5 whole limb buds finding numerous genes involved in embryonic limb morphogenesis and anterior/posterior pattern specification to be differentially expressed (**Fig. 4g-h**). We observed upregulation of the Meis Homeobox 2 (*Meis2)* and *Grem1* genes that are known to be upregulated in the bat FL^50,51^. We also saw the ALX Homeobox 4 (*Alx4*) gene which is a downstream factor of Shh signalling ^52^ to be upregulated in *bZRS^E/E^*. The zinc finger and BTB domain-containing protein 16 (*Zbtb16*, also known as *Plzf*), a transcription factor involved in the development of the anterior-posterior axis in limbs^53^ was also found to be upregulated. *Dlx5;Dlx6* double mutants exhibit hindlimb ectrodactyly ^54^ and both these genes were found to be downregulated in *bZRS^E/E^*. In summary, we found several limb developmental associated genes to be differentially expressed in *bZRS^E/E^* E11.5 embryos.

For the two adjacent *Msx2* BARs (**Fig. 5a**), we first carried out a bat and mouse enhancer assay, cloning both of them together as they were adjacent to each other, finding that the mouse *Msx2* sequence has stronger enhancer activity at E13.5 in the distal autopod while the bat sequence showed reduced expression in this region (**Fig. 5b**, **Extended Data Fig. 6a**). We next swapped this sequence in mice with the bat sequence using the modified PITCh protocol^29^. We found *BAR-Msx2* homozygous mice (*bMsx2^E/E^*) to have significantly reduced expression of *Msx2* in the FL and HL compared to WT mice, with the FL showing a stronger difference (**Fig. 5c**). Analysis of bone ossification using alcian blue/alizarin red staining at E18.5 showed reduced ossification of the 4th intermediate phalanx for *bMsx2^E/E^* compared to wild-type embryos; however, this was not a statistically significant reduction (**Fig. 6a-b**).

**Fig 5:**
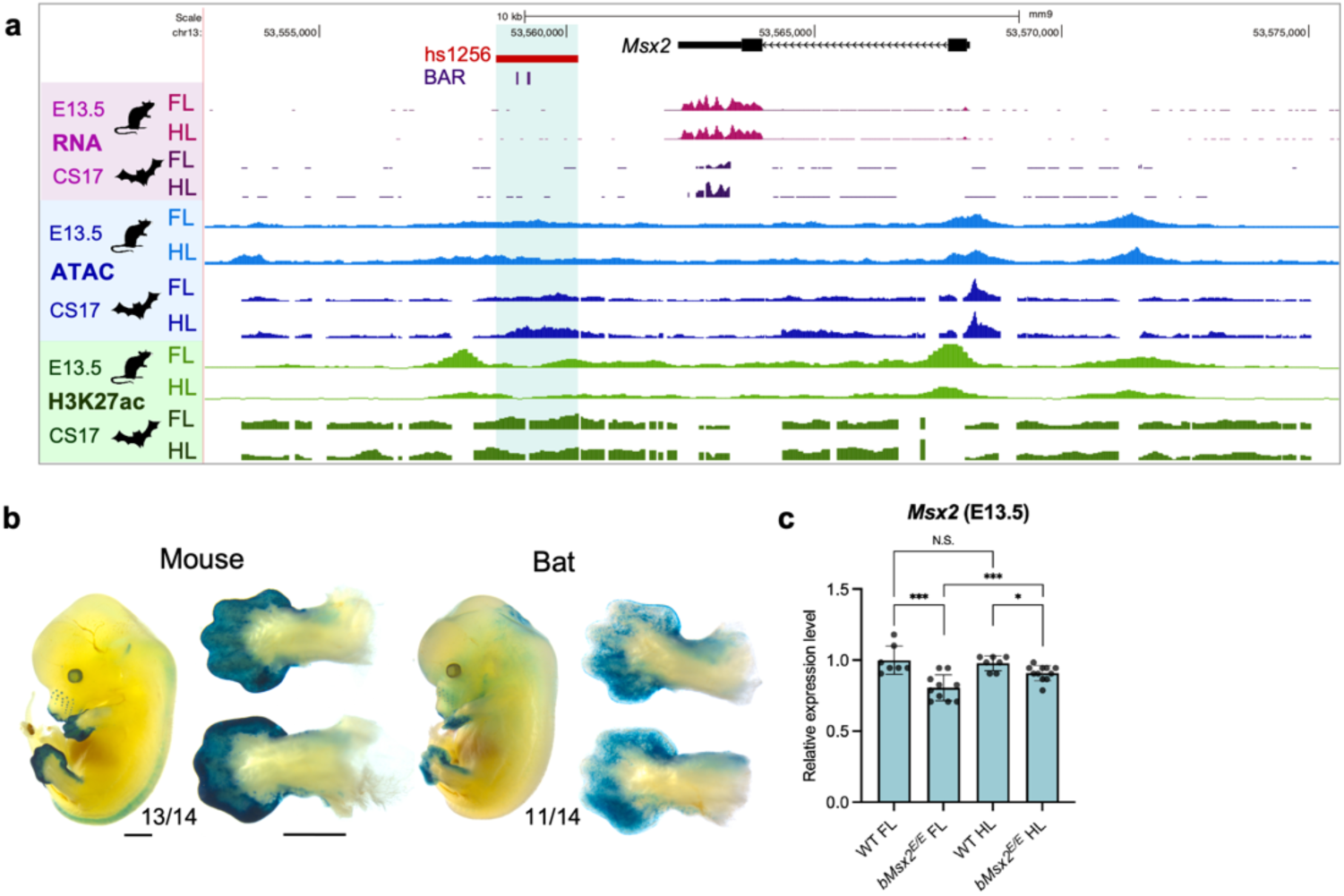
Mouse characterization of the bat *Msx2* enhancer. **a,** Genomic landscape of *Msx2* locus. RNA-seq (pink), ATAC-seq (blue), and H3K27ac ChIP-seq (green) data from bat or mouse FL and HL at E13.5 (mouse) or CS17 (bat). **b,** LacZ enhancer transgenic assay at E13.5 result. The number below the representative embryo picture indicates the number of embryos showing similar enhancer activity. Bars below denote 1mm. **c,** Gene expression levels of *Msx2* in WT and bat enhancer homozygous knockin mice in FL and HL as measured by qRT-PCR. Each value represents the ratio of *Msx2* gene expression to that of *b-Actin* (*n* = 7-11 biological replicates), and values are mean ± standard deviation. The expression value of wild type (WT) FL was arbitrarily set at 1.0. Each dot represents one embryo, and statistical differences were determined using unpaired t test (\*\*\**p* < 0.005, \**p* < 0.05, N.S., not significant).

**Fig 6:**
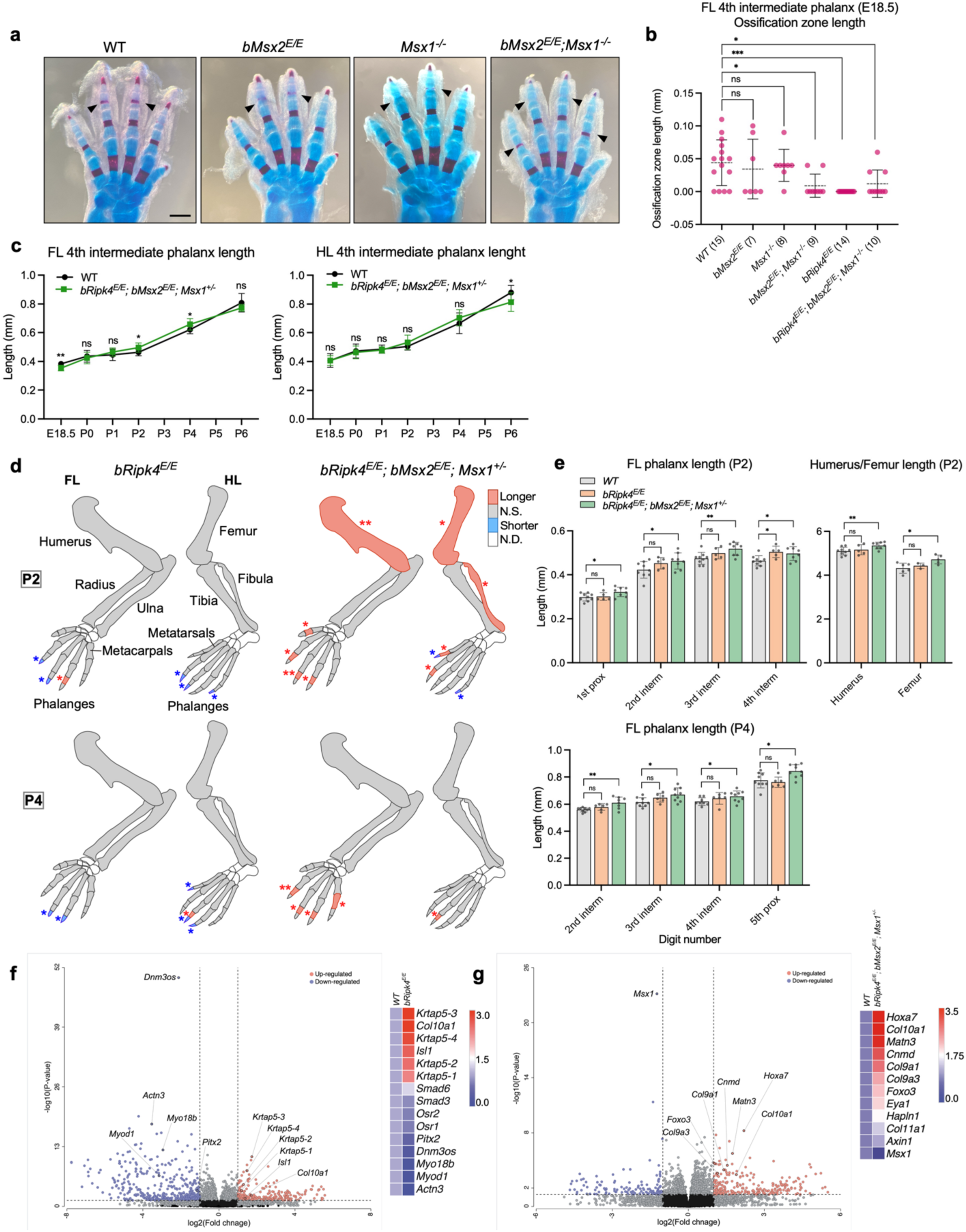
Characterization of compound mice carrying multiple bat enhancers. **a,** Alcian blue and alizarin red staining for WT, *bMsx2^E/E^*, *Msx1^-/-^*, *bMsx2^E/E^; Msx1^-/-^* mice. Bars below denote 0.5mm. Arrowheads indicate regions of delayed ossification. **b,** Ossification zone length of 4th digit intermediate phalanx at E18.5. Each dot represents one embryo. Biological replicate numbers are shown beneath the graph. **c,** Transition of the 4^th^ intermediate phalanx length from E18.5 to P6. **d,** Schematic representation of each bone length difference. Statistical significant P2 and P4 bone length differences between WT and *bRipk4^E/E^* or WT and *bRipk4^E/E^; bMsx2^E/E^; Msx1^+/-^* are shown. **e,** Bone lengths in P2 or P4 WT, *bRipk4^E/E^* and *bRipk4^E/E^; bMsx2^E/E^; Msx1^+/-^.* Each dot represents one embryo (*n* = 5-9 biological replicates). **f** (wild-type vs *bRipk4^E/E^*) **-g** (wild-type vs *bRipk4^E/E^; bMsx2^E/E^; Msx1^+/-^*mice), Left: Volcano plots showing the global transcriptional changes for the indicated groups digits at P4. Each circle represents one gene. The log2 fold change in the indicated genotype is represented on the x axis. The y axis shows the *p* value. A *p* value of 0.05 and a fold change of 2 are indicated by lines. Right: Heat map showing gene expression differences between WT and bat enhancer replacement mice, based on RNA-seq. All statistical differences were determined using an unpaired t test (\**p* < 0.05, \*\**p* < 0.01, \*\*\**p* < 0.005, ns, not significant, nd, not determined).

Limb malformations are only observed when both *Msx1* and *Msx2* genes are altered but not individually^55^, likely due to functional redundancy. Deleting *Msx2* conditionally in the limb using a *Prx1* Cre that drives expression in the limb mesenchyme along with *Msx1* knockout mice showed impairment of the apical ectodermal ridge (AER) and subsequent mesenchyme along with digit abnormalities including preaxial polydactyly^56^. We thus carried out mouse transgenic enhancer assays for five putative *cis*-regulatory elements bat sequences spread broadly over 100 kb of intergenic regions in the bat *Msx1* locus (**Extended Data Fig. 7a**). We did not observe any differences in activity between FL and HL (**Extended Data Fig. 7b**). We thus set out to test whether placing *bMsx2^E/E^* mice on an *Msx1* heterozygous gene knockout background instead of enhancer replacement will exacerbate the limb phenotype. Using the *i-*GONAD protocol^57^, we generated *Msx1* heterozygous gene knockout mice (*Msx1^+/-^*). *Msx1* homozygous mice are known to die at birth^58^. We then crossed *Msx1^+/-^* gene knockout mice to *bMsx2^E/E^* mice obtaining *bMsx2^E/E^; Msx1^-/-^* compound mice. As *Msx1* homozygous knockout mice die at birth, we carried out alcian blue/alizarin red staining at E18.5 finding these mice to have significantly reduced ossification in the 4^th^ intermediate phalanx compared to *bMsx2^E/E^* or *Msx1^-/-^* mice (**Fig. 6a-b**). These results suggest that lower *Msx1* expression levels leads to reduced compensation for differential expression of *Msx2*, leading to a stronger limb ossification phenotype.

### Compound mice show bat associated phenotypes

We next set out to test whether combining bat-mouse swapped alleles could have an additive effect on phenotype. As both *bRipk4^E/E^*and *bMsx2^E/E^; Msx1^-/-^* mice showed delayed bone ossification in the digits, we mated these lines to one another. We observed that *bRipk4^E/E^; bMsx2^E/E^; Msx1^-/-^* mice (triple compound mice) had significantly reduced ossification at E18.5 (**Fig. 6a-b**). In bats, delayed ossification in the FL was observed as a means to allow for longer digit length^2,3^. Additionally, *Msx2* knockout mice showed defective chondrogenesis and endochondral ossification^59,60^. We thus examined the length of the 4th intermediate phalanx in WT and triple compound mice FL and HL from E18.5 to P6 using alcian blue and alizarin red staining of bone specimens, as this is one of the bones showing significant differences during the neonatal period. As *Msx1* homozygous knockout mice are lethal at birth we used *Msx1* heterozygous mice (*Msx1^+/-^*) for these analyses. Forelimb bone length was comparable or slightly shorter from E18.5 to P1, increased at P2 and P4, and showed no difference by P6 (**Fig. 6c**). The HL showed a similar trend, but no significant difference was observed from E18.5 to P4 (**Fig. 6c**). We measured all bone lengths at P2 and P4 in *bRipk4^E/E^* and triple compound mice, and compared with WT (**Fig. 6d**). Although both *bRipk4^E/E^* and triple compound mice showed delayed bone ossification in the digits, bone length differences were more pronounced in the triple compound mice (**Fig. 6d-e**). *bRipk4^E/E^* mice had shorter distal phalanges compared to WT at P2 and P4. In contrast, triple compound mice showed increased bone length at P2 in multiple bones, including the phalanges, humerus, femur, and fibula (**Fig. 6d-e**). Four forelimb phalanges and one hindlimb phalanx remained longer at P4 (**Fig. 6d-e**).

To identify genes involved in these differences, RNA-seq was carried out on wild-type, *bRipk4^E/E^* and *bRipk4^E/E^; bMsx2^E/E^; Msx1^+/-^* mice from P4 digits (**Fig. 6f-g**). In *bRipk4^E/E^*, we observed upregulation of *Krtap 5-1, 2, 3* and *4* genes that are major components of hair and nails^61,62^ (**Fig. 6f**). We also saw downregulation of ossification related genes, such as *Osr1* and *Osr2*. Dominant negative *Osr2* transgenic mice, in which the function of both *Osr1* and *Osr2* genes is inhibited, show delayed mineralization in calvarial and tibial bone tissues^63^. *Smad3* and *Samd6* were slightly down- and up-regulated respectively. TGF-β/Smad3 signals are known to repress chondrocyte hypertrophic differentiation^64^ and overexpression of *Smad6* significantly delays chondrocyte hypertrophy, which can lead to postnatal dwarfism with osteopenia^65^. *Col10a1* is a marker gene of hypertrophic chondrocytes^66,67^ that was also found to be upregulated.

These results suggest there are perturbations of the endochondral ossification process in digits.

In the triple compound mice, we observed upregulation of skeletal development-related genes, including *Hoxa7*, *Cnmd, Foxo3* and *Eya1*, as well as extracellular matrix (ECM) genes enriched in growth plate chondrocytes, such as *Col9a1*, *Col9a3, Col10a1* and *Matn3*^67,68^, along with structural and matrix-organizing ECM components *Col11a1* and *Hapln1*(**Fig. 6g**). In the context of endochondral ossification, *Cnmd* (ChM-1; Chondromodulin-I) is a negative regulator of cartilage vascularization, maintaining avascular cartilage and thereby controlling the timing of ossification^69^. *Foxo3* inhibits Wnt signaling and delays ossification^70^. Overall, our transcriptomic analyses show inhibited Wnt signaling and delayed ossification, marked by retained hypertrophic chondrocytes, upregulation of collagen genes, and a potential role in prolonged bone elongation in triple compound mice.

## Discussion

The bat wing is a unique organ that underwent numerous morphological changes during evolution to allow for powered flight. Here, we used bat and mouse embryos to identify in a genome-wide manner numerous genes and regulatory elements that show unique activity in the developing bat FL. Using mouse transgenic enhancer assays, we identified several sequences that show differential limb enhancer activity between the bat and mouse sequence. We then swapped six sequences in the mouse genome with the corresponding bat sequence finding relevant morphological changes associated with both growth and patterning aspects of the bat limb phenotype, including delayed bone ossification, slightly longer digits, increased epidermal layer thickness and symmetrical hindlimbs. Combining swapped lines that affected bone ossification had a small but significant additive effect on the phenotype within a specific neonatal time window and led to longer phalanges, humerus, femur, and fibula.

Our genomic datasets (RNA-seq, ChIP-seq, and ATAC-seq) provide useful catalogs for bat limb development. By generating matching developmental time points for mouse FL and HL these datasets can be scanned for differential gene expression and regulation across these two species. We focused our analyses on differential activity of homologous sequences between the mouse and bat genome. However, in the future it will be intriguing to also characterize sequences that are either unique to bats or missing in bats but appear in other mammals, which could also be associated with limb development. In addition, further analyses of noncoding RNAs, such as miRNAs and lncRNAs, that are associated with bat wing development would also be of interest.

The bat *Ripk4* regulatory sequence led to the strongest phenotypic effect from an individual swapped sequence. This included both a delay in bone ossification, mild changes in bone length and a thicker epidermal layer. Wnt signaling has been shown to play an important role in limb organogenesis^71^. Our previous bat FL and HL genomic characterization^13^ showed an important role for this pathway in wing development with higher levels of canonical Wnt pathway antagonists and an overall suppression of the canonical Wnt/β-catenin pathway in the bat FL compared to HL and an increased expression of canonical Wnt receptors in the bat HL^13^. Ripk4 phosphorylates Dishevelled (Dvl) which leads to an increase in β-catenin accumulation and promotes the activation of the canonical Wnt signaling^72^. We observed increased activity for the bat *Ripk4* enhancer in the developing digits of both FL and HL, likely leading to increased Wnt activity. Mouse knockouts for *Ripk4* show shorter limbs, fusion of digits and reduced skin folds^31^. Thus, increased activity of *Ripk4* and subsequently the Wnt pathway will likely lead to an opposite phenotype of longer limbs and thicker skin folds, as we observed in *bRipk4^E/E^* mice. While we characterized the limb bud, *Ripk4* has also been shown to be highly expressed in the bat patagium^33^ and could also have an important role in shaping this structure. Our genomic data showed additional regions to be differentially active between bats and mice in this locus (**Fig. 3a**), suggesting that additional regulatory elements might be involved in various wing associated phenotypes.

The triple combined line, *bRipk4^E/E^; bMsx2^E/E^; Msx1^+/-^* had small additive effects on phenotype with mice exhibiting a more pronounced phenotype than *bRipk4^E/E^* by itself at P2 and P4. These included delayed bone ossification in the digits and increased bone length including the phalanges, humerus, femur, and fibula (**Fig. 6d-e**). Bats are known to have delayed ossification in the forelimb^3^. RNA-seq analyses of these triple compound mice showed increased inhibition of the Wnt pathway, which is known to contribute to delayed ossification^73,74^. Endochondral ossification is a key process in longitudinal bone elongation, characterized by an increase in chondrocyte number and size, followed by matrix calcification and subsequent replacement of cartilage with bone, ultimately forming trabecular bone. Our results suggest that multiple regulatory factors involved in the attenuation of Wnt signaling might be responsible for delayed ossification in bats. In terms of longitudinal bone elongation, not only delayed ossification but also increasing both the size and number of chondrocytes are essential^75,76^. Further work could explore the factors involved in chondrocyte phenotype to drive bone elongation.

*Msx2* is known to be involved in endochondral ossification and stimulate the maturation of chondrocytes into the prehypertrophic and hypertrophic stages^59,60^. However, in our study, we focused on earlier developmental stages, and indeed, our *bMsx2^E/E^* mice showed gene expression changes at E13.5 but not at P4, during the endochondral ossification stage. This suggests that other putative regulatory elements may be involved in the stage-specific regulation of *Msx2*. Based on our genomics dataset, we did not detect differential regulatory regions associated with key limb development genes such as *Fgf*, *Bmp*, *Pthlh*, *Sox9*, or *Ihh*. There are several potential reasons for this: 1) Our targeted developmental stages may be too early to capture *Pthlh*, *Sox9*, or *Ihh* activity; 2) We used bulk RNA-seq, which may not identify expression changes in smaller cell populations, such as localized expression of *Fgf8* or *Shh*. It is also worth noting that a recent study that carried out single-cell RNA-seq on developing mouse and bat (*Carollia perspicillata*) FL and HL at E11.5, E12.5 and E13.5 in mice and CS15 and CS17 in bats did not observe differential expression in *Fgf* or *Bmp*^51^. From our results we did have an observable limb phenotype in *bZRS^E/E^* mice, while this sequence did not show any significant differences in our genomics data; and 3) gene expression changes during early development may lead to phenotypic differences at later stages. For example, in the triple compound mice, we did not detect differential expression of *Msx2* or *Ripk4* at P4, despite clear differences in ossification and bone length at that stage. This highlights the complexity and importance of capturing dynamic gene regulation within specific developmental time windows, which remain a challenge but also an opportunity for future temporal-resolution genomics.

Interestingly, swapping the mouse sequence with a bat accelerated region, BAR116, that showed strong FL enhancer activity for the bat sequence^17^ and chicken sequence^43^ but was negative for the mouse sequence did not lead to an observable phenotype. The *HoxD* locus contains a cluster of nine *HoxD* genes that are regulated in a spatiotemporal manner by multiple enhancers whose chromatin interactions are thought to be regulated by two distinct topologically associating domains (TADs). When a distal *Hoxd* limb enhancer is inserted into the proximal limb TAD, its distal limb-specific activity is suppressed in this ectopic context^77^, suggesting that the native chromatin environment can mask the activity of exogenously inserted enhancers even in the same *Hoxd* locus. To overcome this limitation and more accurately recapitulate endogenous *cis*-regulatory function in bats, strategies such as TAD-level structural reconstruction or the use of longer transgenes (e.g., BAC/YAC constructs) might be needed.

For the ZRS, we primarily obtained a bat-HL phenotype. This sequence was chosen due to it being a BAR. However, we did not observe strong differences between bat and mouse ZRS enhancer activity and also between bat FL and HL in our genomic analyses (**Extended Data Fig. 8a**). However, we reasoned that due to the strong phenotypic effects of point mutations in this enhancer that lead to a multitude of limb developmental phenotypes in humans, mice and other mammals^46,78,79^, we would likely observe a limb development associated phenotype. Interestingly, while we did not obtain a strongly penetrant FL-associated phenotype, we did observe a HL phenotype in *bZRS* mice that included polydactyly and the formation of symmetrical HL digits.

A key region that differs between bats and mice in the ZRS is a known binding site for the ETS translocation variants 4 and 5 (Etv4/5) proteins. Posterior *Shh* gene expression in the limb bud is regulated by a balance between transcriptional activators of the E26 transformation-specific (ETS) family and the repressors Etv4/5^80^. The ZRS contains multiple binding sites for both ETS and ETV factors (**Extended Data Fig. 8b**). *ETS1/2* are predominantly expressed in the posterior region of the limb bud, while *Etv4/5* are expressed throughout the distal limb bud, encompassing both anterior and posterior regions. The spatial expression patterns of these factors define the posterior-specific expression of *Shh*. Mutations in ETV binding sites can disrupt this balance, leading to ectopic activation of *Shh* in the anterior limb bud^80^. For example, conditional knockout of *Etv5* in mesoderm-derived cells using *Brachyury* (*T*)-*Cre* leads to anterior reactivation of *Shh* and results in polydactyly specifically in the hindlimbs, mirroring the phenotype observed in *bZRS* mice^81^. In the *bZRS* sequence, we identified sequence differences within the ETV-A site and its surrounding region, which overlap with variants previously linked to triphalangeal thumb polydactyly^82–85^ (**Extended Data Fig. 8b-c**). Notably, the rs606231152 variant (295T>C, represented as A>G in the figure due to reverse complement orientation) is highly dominant over the wild-type allele in families with triphalangeal thumb polydactyly. Intriguingly, most bat species examined also carry the A>G substitution at this site (**Extended Data Fig. 8b-c**). Testing the human ZRS sequence with the rs606231152 A>G change in a mouse enhancer assay showed that it leads to anterior enhancer activity^85^. In addition, it was shown that mutating the ETV-A site alone is insufficient to drive anterior enhancer activity in a LacZ transgenic assay^80^, suggesting that multiple mutations within the ZRS, including variants like rs606231152, are likely required to induce robust anterior enhancer function. Interestingly, a recent study highlighted the importance of a low-affinity ETS binding site (ETS-A)^86^, which we identified as partially overlapping with the aforementioned ETV-A site^80^ (**Extended Data Fig. 8b**). High-affinity mutations in ETS-A lead to anterior reactivation of *Shh* and result in a similar hindlimb polydactyly phenotype as observed in our *bZRS* mice. It is possible that ETS binding to ETS-A interferes with ETV-A binding, suggesting that competitive binding and the balance between ETS and ETV factors, particularly in overlapping ETS-A/ETV-A sites, could play a crucial role in this phenotype.

Our results suggest that the bat wing arose from numerous small molecular changes, in line with past work showing that morphological novelty in evolution likely arose from numerous changes^23^. In bats, comparative evolutionary genomics is particularly difficult as there are no intermediate species from the fossil records^87^ thus restricting the ability to decipher the morphological skeletal evolution that leads to bats. Recent work comparing bird and bat limb evolution by analyzing hundreds of skeletal structures suggests that the bat FL and HL evolved in unison in contrast to birds where they evolved independently thus restricting each other’s development and morphology^6^.

Thus, it could be that analyzing changes driving both FL and HL phenotypes could identify sequences that are more likely to drive wing development. As mentioned above, our top phenotype altering sequence, *bRipk4*, showed increased enhancer activity in both FL and HL.

Our study had several limitations. We focused our genomics work on three bat developmental time points, thus potentially missing earlier or later time points that could also be important in wing development. We selected sequences primarily in a biased manner, focusing on those that are thought to regulate known genes involved in limb development and malformations. We only changed the targeted mouse genomic sequences to one specific bat species, *M. natalensis*. Other bat sequences could have a different effect, due to the large diversity of wing morphology in bats. To circumvent this, we could have predicted the last common ancestor sequence of bats and swapped this sequence in, but it may not have represented a current existing bat. As bat genome editing is currently not possible, we modeled the hypothesised genomic changes in mice whose chromatin architecture or ‘trans’ environment could have a differential effect on limb development and might not be a good model to assay for limb elongation, as to our knowledge, there are very limited mouse mutants that have an elongated limb phenotype. Finally, we focused our work on individual alleles and only generated one combined allele.

In summary, our work provides a detailed genomic catalog of candidate genes and putative *cis*-regulatory elements involved in bat FL and HL development, compared with corresponding stages in mice. Through extensive *in vivo* functional analyses, we demonstrate the phenotypic relevance of several of these elements, highlighting the role of enhancers in bat wing morphogenesis and suggesting that multiple regulatory changes contributed to the emergence of the bat wing, one of the most eminent evolutionary adaptations in mammals.

## Methods

### RNA-seq

RNA was extracted from paired forelimbs and hindlimbs from three C57BL/6J (Jackson labs, 000664) embryos (biological replicates) at three developmental stages (E12.0, E13.0, E13.5) using the RNeasy Midi (Qiagen) kit. Total RNA samples were enriched for poly-A containing transcripts using the Oligotex mRNA Mini kit (Qiagen) and strand-specific RNA-seq libraries were generated using PrepX RNA library preparation kits (IntegenX) following the manufacturer’s protocol. After clean up with AMPure XP beads (Beckman Coulter) and amplification with Phusion High-Fidelity polymerase (NEB), RNA libraries were sequenced on a HiSeq 2500 to a depth of at least 30 M reads. All mouse work (other than ATAC-seq which was approved by the University of Cape Town) was approved by the UCSF Institutional Animal Care and Use Committee (IACUC) protocol number AN197608. Sequence alignment was performed using STAR^88^. Mappings were restricted to those that were uniquely assigned to the mouse genome; unique read alignments were used to quantify expression and aggregated on a per-gene basis using the Ensembl GRCm37.67 annotation. We analysed these raw data with DESeq2^89^ to assess variance and differential expression.

### ChIP-seq

C57BL/6J (Jackson labs, 000664) mouse forelimbs and hindlimbs dissected from E12.0, E13.0, E13.5 were cross-linked with 1% formaldehyde for 10 minutes, quenched with glycine and flash frozen. Cross-linked limbs were then combined into pools of 4-7 pairs per stage for chromatin sheering using a Covaris S2 sonicator. Sheared chromatin was then used for chromatin immunoprecipitation (ChIP) with antibodies against active (anti-H3K27ac; Abcam ab4729) or repressed (anti-H3K27me3; Millipore 07-449) chromatin marks using the Diagenode LowCell# ChIP kit following the manufacture’s protocol. Libraries were prepared using the Rubicon ThruPLEX-FD Prep Kit following the manufacturer’s protocol and sequenced on an Illumina HiSeq2500 using single end 50 bp reads to a sequencing depth of at least 25 M. Raw sequence files were aligned from fastq to mm9 or mnat_v1 bam format using samtools^90^ and a pre-built annotation index of the respective assembly. ChIP-Seq files include an input channel for each species, stage, and tissue. This allowed us to obtain increased accuracy when calling the peaks. Peak calling was done two-fold. First, using MACS v2^91,92^ we called the peaks on each of the channels separately, then with elaborate background processing. In the first step, MACS^91^, we used the following parameters: *macs2 -n file-name -t file-name.sort.bam -c input-file.sort.bam -g mm --nomodel --pvalue=5e-5 --slocal=5000 --llocal=20000 --shiftsize=300* for H3K27ac. We used *macs2 -n file-name -t file-name.sort.bam -c input-file.sort.bam -g mm --nomodel --pvalue=1e-4 --slocal=10000 --llocal=50000 --shiftsize=500* for H3K27me3. In the bat we used the same parameters for either antibody: *--nomodel --pvalue=1e-5 --slocal=5000 --llocal=20000 --shiftsize=30*0. Later, we added background processing from^93^ to improve the ability to align the called peaks across the channels. For example, peaks called on E12.0FL and E12.0HL are expected to misalign. Using the background analysis of Diaz et al and the background profiling provided by deepTools^94^, we were able to orient and identify the same called peaks. After peak calling for each channel we aligned the peaks across the species. Misaligned peaks could be missing from the other species but as we do not having any prior information about the nature of these regions, we did not use them in a differential analysis context. To characterize regions across the species, we used lifOver^95,96^ allowing multiple targets to be listed in the output. After obtaining all the lifted-over targets, we employed a search algorithm for synteny anchors on either side of each peak. To match the best candidate peak across the two species, the algorithm was required to observe two synteny blocks in the flanks of each peak (see **Extended Data Fig. 9**). We then used the following statistical analysis to list the differential chromatin (Differentially Active) regions across bat and mouse. We created a design that accounts for chromatin activity levels in all the possible combinations of species (2 - mouse vs. bat), tissue (2 - FL vs. HL), and stage (3 - E12.0, E13.0, or E13.5). We used the R package DESeq2^89^, which we customized to use counts estimated from FPKM using the normalization method due to^93^, then used these numbers only on aligned regions. The count data were averaged across the replicates, and we then performed Maximum Likelihood Estimation (MLE) given two models; a simple model where the main effects considered are *Species + Stage*, and an elaborate model whose main effects are further enhanced by interactions: *Species + Stage + Tissue + Tissue:Stage + Tissue:Species*. The MLE between the two models that was marked after FDR-correction as statistically significant, gave the differential chromatin (DA) regions listed in the supplementary master_features_v3_pvalSort_20230806.xlsx available in Zenodo.

### ATAC-seq

Mouse embryos for ATAC-seq (C57BL/6 strain UCT3) were supplied by the Animal Research Facility, University of Cape Town. Ethical approval for their use was granted by the Health Science Animal Ethics Committee (FHS AEC014/07). *Miniopterus natalensis* embryos were collected from wild-caught, pregnant females in 2016 and 2017 from De Hoop Nature Reserve, Western Cape Province, South Africa (Western Cape Nature Conservation Board permit number: AAA007-00133-0056). Approval for the use of the bat embryos was granted by the Science Faculty Animal Ethics Committee (2012/V41/NI and 2012/V39/NI). Limbs were dissected from embryos and forelimbs and hindlimbs from the same embryo were pooled separately and placed in solutions of 0.25%Trypsin/0.1M EDTA:HBSS at 1:1 or 1:3 dilution and incubated at 4 °C overnight. For all E13.5 mouse limbs, limbs from 4 embryos were pooled. Samples were centrifuged at 250 x *g* for 3 minutes, all but 1 ml of solution removed, and incubated at 37 °C for 25 minutes. A further 1 ml of 0.25% trypsin/0.1M EDTA was added, followed by incubation at 37 °C for 100 minutes. Limbs were dissociated by gentle pipetting and centrifuged at 250 x *g* for 5 minutes to pellet the cells. The solution was removed and pellets were washed in Hanks followed by a further centrifugation at 250 x *g* for 5 minutes. Cells were resuspended in DMEM with 10% fetal calf serum and quantified using a haemocytometer. 50,000 cells were centrifuged at 500 x *g* for 5 minutes at 4 °C, washed twice with ice cold 1x PBS and centrifuged for a further 5 minutes. The cell pellet was resuspended in cold lysis buffer (10 mM Tris-HCl, pH7.4; 10 mM NaCl; 3 mM MgCl2; 0.1% IGEPAL CA-630), and centrifuged for a further 10 minutes at 500 x *g* at 4 °C. The supernatant was discarded and the cell nuclei resuspended in a transposition reaction mix [25 μl 2x TD Buffer (C-121-1030, Illumina); 2.5 μl Tn5 Transposase (FC-121-1030, Illumina); 22.5 μl Nuclease Free H2O]. The reaction was incubated at 37 °C for 30 minutes. To stop tagmentation, EDTA was added to a final concentration of 50 nM and the reaction was incubated at 50 °C for 30 minutes. Tagmented DNA was cleaned up using Qiagen PCR purification MinElute Kit (28004, Qiagen) and amplified using KAPA HiFi Hotstart ReadyMix (TAQHSKB, Roche) for 12-14 cycles. The amplified DNA was purified using a Qiagen PCR purification MinElute kit (28004), and KAPA Pure Beads (KK8000, Roche). Tagmentation efficiency was assessed using Agilent BioAnalyzer and libraries quantified using a Qubit Fluorometer (Thermo Fisher) at the Central Analytical Facility (University of Stellenbosch). The concentration of ATAC library pools was assessed with the KAPA Library Quantification Kit (KK4835, Roche).

CS17 bat and E13.5 mouse libraries were sequenced by BGI (Hong Kong) using paired-end 50 bp runs on the Illumina HiSeq4000 platform. We used a three-stage MACS v2^91,92^ to eliminate background. First, we called peaks assuming a flat model: *macs2 callpeak -t bedGraph-file-name -B --nomodel --pvalue=5e-5 --slocal=5000 --llocal=20000 --SPMR -n file-name*. We then performed an enrichment estimation by creating a treatment vs. control statistical apparatus: *macs2 bdgcmp -t treatment_pileup.bdg -c control_lambda.bdg -o FE.bdg -m FE*. We then subtracted the noise residual of the two models: *macs2 bdgcmp -t treatment_pileup.bdg -c control_lambda.bdg -o sub.bdg -m subtract*. The ATAC-Seq channels were utilized in order to home in on the exact locations of the chromatin action in regions of differential chromatin (DA).

### LacZ enhancer transgenic assays

Mouse work was approved by the UCSF Institutional Animal Care and Use Committee (IACUC), protocol number AN197608, and was conducted in accordance with AALAC and NIH guidelines. The bat and mouse enhancer regions were amplified from mouse or bat (*Miniopterus natalensis*) genomic DNA by PCR, cloned into the Hsp68-LacZ vector^97^ and sequence verified. Cloned regiones are shown in **Data S2**. All LacZ transgenic mice were generated by Cyagen Biosciences using standard procedures^98^, and harvested and stained for LacZ expression as previously described^99^ at specific time points noted on figures. Pictures were obtained using a M165FC stereo microscope (Leica).

### Generation of bat enhancer replacement mice

We used the Precise Integration into Target Chromosome (PITCh) system^29^ with minor modifications. Briefly, homology between bat and mouse sequences was characterized using CLUSTALW^100^. For donor plasmid construction, the bat enhancer and 1 kb mouse homology arms (5′ and 3′) were amplified by PCR. The cloned regions are listed in **Data S2.** These three fragments were cloned into the backbone plasmid pLSODN-3 (Biodynamics Laboratory, DS620) or pGEM-T easy (Promega, A1360) using NEBuilder HiFi DNA Assembly Master Mix (New England Biolabs, E2621L). Circular donor plasmids were dialyzed with a membrane (Millipore, VSWP04700) and used for pronuclear injection. Two gRNAs were designed to target the 5′ and 3′ ends of the mouse endogenous homology region (light blue in **Fig. 2d**) using the gRNA design tool on the Integrated DNA Technologies (IDT) website and were selected based on low off-target and high on-target scores. One of the gRNA target sites was placed at the 5′ and 3′ end of the homology in the donor plasmid (green in **Fig. 2d**). gRNAs are listed in **Supplementary Table 1**. A mixture of crRNA (IDT), tracrRNA (IDT), Cas9 protein (IDT), and donor plasmid was used for pronuclear injection. FVB/NJ mouse stain (Jackson Laboratory; 001800) was used for all bat enhancer replacement mice generation. Injections were performed at the Gladstone Transgenic Core. Founder mice were screened by PCR and sequencing.

### Alcian blue and alizarin red staining

Embryos or neonate mice limbs were harvested at E18.5 and limbs were dissected and used for staining. Alcian blue/Alizarin red staining was performed according to standard procedures for late-gestation stage embryos^101^. Briefly, following skin removal, limb tissues were fixed in 95% ethanol overnight at room temperature, then treated with acetone overnight at room temperature. Samples were stained with Alcian blue overnight at room temperature and destained with sequential quick washes in 70% and 95% ethanol, followed by overnight incubation at room temperature. Tissues were then pre-cleared in 1% KOH for 1 hour at room temperature, stained with Alizarin red for 3–4 hours at room temperature, and incubated in a 1:1 solution of 50% glycerol and 1% KOH to remove excess red stain. Ossification zone and phalanx bone lengths were measured using M165FC stereo microscope (Leica).

### Histology

Whole limb tissues were fixed in 4% PFA at 4 °C overnight. The dorsal and ventral skin from the arm was then dissected and further fixed in 4% PFA for 3 hours. Skin tissues were embedded in paraffin, sectioned at 10 μm using a rotary microtome, and used for standard Hematoxylin and Eosin (H&E) staining^102^. Epidermis thickness was measured using Fiji software^103^.

### RNA extraction and qPCR

Total RNA was collected from tissue using TRIzol (Thermo Fisher Scientific; 15596026) and converted to cDNA using ReverTra Ace qPCR-RT master mix with genomic DNA (gDNA) remover (Toyobo; FSQ-301). qPCR was performed using SsoFast EvaGreen supermix (Bio Rad; 1725205). Primer sequences used for qPCR are shown in **Supplementary Table 2**.

### RNA-seq for bat enhancer replacement mice

RNA-seq was conducted by Novogene. The library was generated with NEBNext Ultra II RNA Library Prep Kit for Illumina (NEB; 7770), and sequencing was done using the NovaSeq 6000 S4 platform with PE150. The data were analyzed using Partek Flow (Version 10.0). After primary quality assessment was performed, bases and reads with low quality were filtered out and the reads were aligned to the mouse reference genome (mm10) using STAR (Version 2.7.8a)^88^. The final BAM files were quantified using the Partek E/M algorithm^104^. Normalization of read count was performed by the total number of counts (count per million) plus 0.0001, and all genes with less than ten normalized read counts were excluded from subsequent analyses. Differentially expressed genes were identified using DEseq2^89^ in Partek Flow. Gene ontology^37^ and Morpheus (RRID: SCR_017386, https://software.broadinstitute.org/morpheus) were used for data analysis.

### Whole-mount *in situ* hybridization

For bat embryos, whole-mount in situ hybridization was carried out as described in ^8^. For mouse, E11.5 embryos were fixed in 4% paraformaldehyde. A plasmid containing mouse *Shh* and *Ptch1* cDNA (GenScript; Shh_OMu22903D and Ptch1_OMu22749D) were used as templates for DIG-labeled probes. Mouse whole-mount in situ hybridization was performed using standard procedures^105^.

## Acknowledgements

We want to thank Dr. Hai P. Nguyen for discussion, advice and encouragement and the Ahituv lab for helpful advice. This work was funded in part by the National Human Genome Research Institute grant numbers R01HG012396 (NA) and K99HG012576 (AU) and R00HG012576 (AU), Japan Society for the Promotion of Science (JSPS) postdoctoral fellowships for research abroad (AU), Uehara Memorial Foundation postdoctoral fellowship (AU) and the Sydney Brenner Fellowship from the Academy of Science of South Africa (DH). US-Israel Binational Science Foundation (no. 2015307, NA and TK) and the Bergmann Memorial Research Award (TK).

## Author information

Contributions

Conceptualization: AU, GK, DH, NI, TK, NA

Methodology: AU, GK, RS, EM, WE, YZ, MN, RR, KF, VB, KN, SK, SAS, MM, SZ, ML, SF, VS, TS, DH

Investigation: AU, GK, RS, EM, YZ, MN, RR, KF, VB, KN, SK, SAS, MM, SZ, ML, SF, MYH, VS, TS, DH, NI, TK, NA

Visualization: AU, GK, RS, EM, MN, KF, VB, SZ, ML, MYH, DH, NI, TK, NA

Funding acquisition: AU, DH, NI, TK, NA

Project administration: AU, GK, DH, NI, TK, NA

Supervision: AU, NI, TK, NA

Writing – original draft: AU, GK, DH, NI, TK, NA

Writing – review & editing: AU, GK, DH, NI, TK, NA

## Supplementary information

Extended Data Figures 1-9

Supplementary Tables 1-2

Supplementary Data 1-2: Excel file containing additional data too large to fit in a PDF

