## Supplementary Material for "Functional characterization of bat limb regulatory elements"

**a**

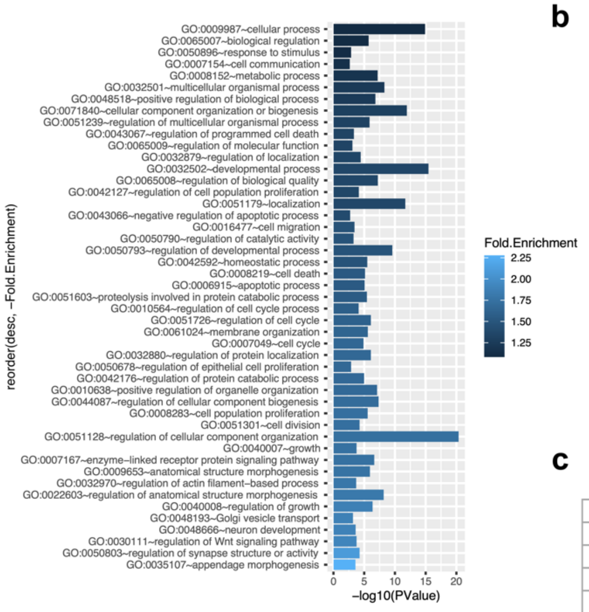


**b**

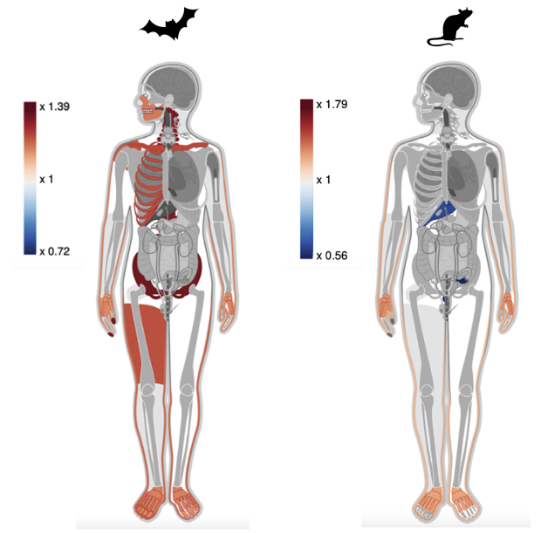


**c**

| | | | | | |
|---|---|---|---|---|---|
| *Adam* | *Atl1* | *Atp6v1b2* | *Atr* | *B9d1* | *Bmpr1b* |
| *Cav1* | *Cep120* | *Cep290* | *Cers3* | *Col11a1* | *Col7a1* |
| *Ctsc* | *Cyp7b1* | *Eln* | *Ext1* | *Fat4* | *Fuca1* |
| *Grip1* | *Hpgd* | *Ift80* | *Inpp5e* | *Insr* | *Itgb4* |
| *Jak2* | *Kctd1* | *Kif1a* | *Kras* | *Lmbr1* | *Lmx1b* |
| *Lrp4* | *Mbtps2* | *Msx1* | *Nfkb1* | *Orc4* | *Pam* |
| *Pde4d* | *Pdgfrl* | *Piga* | *Plec* | *Pvrl1* | *Rab3gap1* |
| *Rfxank* | *Slc40a1* | *Smarcad1* | *Smarcb1* | *Sos1* | *Sos2* |
| *Stat3* | *Stk11* | *Tcf4* | *Tfap2a* | *Tmem231* | *Ube2a* |
| *Wnk1* | *Wnt5a* | *Xylt1* | *Zmpste24* | | |

**d**

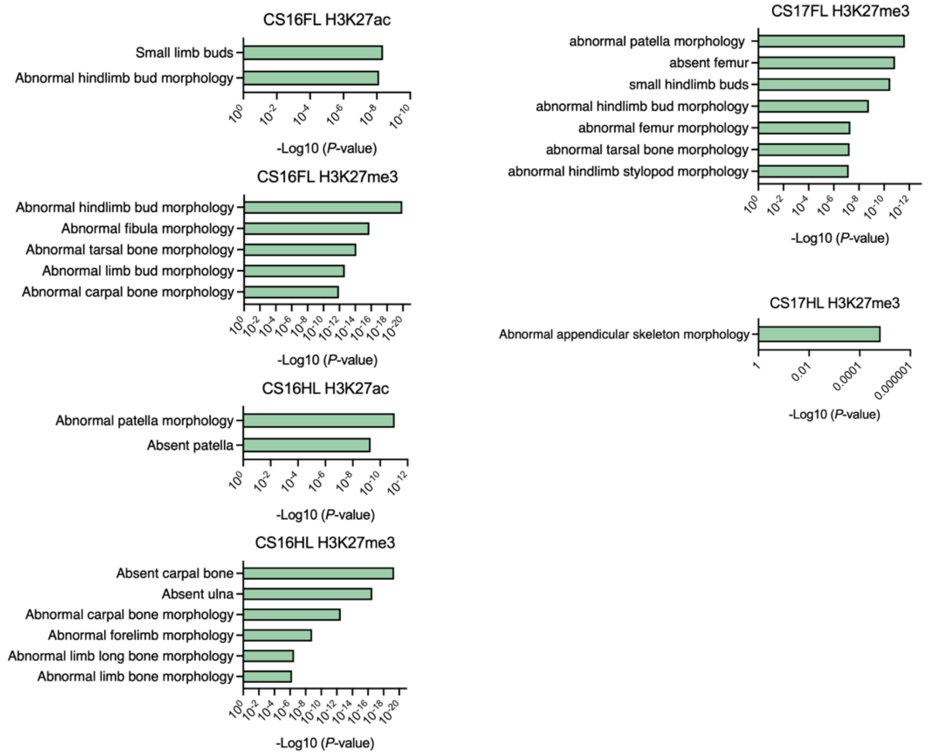


**Extended Data Fig. 1: Gene enrichment analyses for bat versus mouse genomic datasets**
**a,** Gene ontology analysis using DAVID(*20*) of bat-specific DE genes. In the GO biological process analysis of the 1234 bat-DE genes there are 1160 GO terms, out of which 355 are statistically significant

(FDR corrected P-val < 0.05). We pruned the 355 GO terms using Revigo(*90*) into 47 "parent" terms. Inside these 47 groups we estimated the median fold change (color), and p-value (bar size).
**b,** GeneORGANizer(*21*) view of organs that have differentially expressed (DE) genes in bat (left) or mouse (right). Enrichment or depletion are marked with warm (red) vs. cold (blue) colors. The intensities of the colors depicted in the legend are enrichment levels (or odds ratios).
**c,**Table showing DE genes that are enriched in the fingernails out of the enrichment analysis for bat-specific DE genes.
**d,** GREAT(*22*) analysis of DA H3K27Ac and H3K27me3 ChIP-seq peaks between bat and mouse FL and HL compared to the read count pattern of the corresponding region in the mouse. No limb-related associated terms were found for CS17 H3K27ac FL and HL and hence not shown.

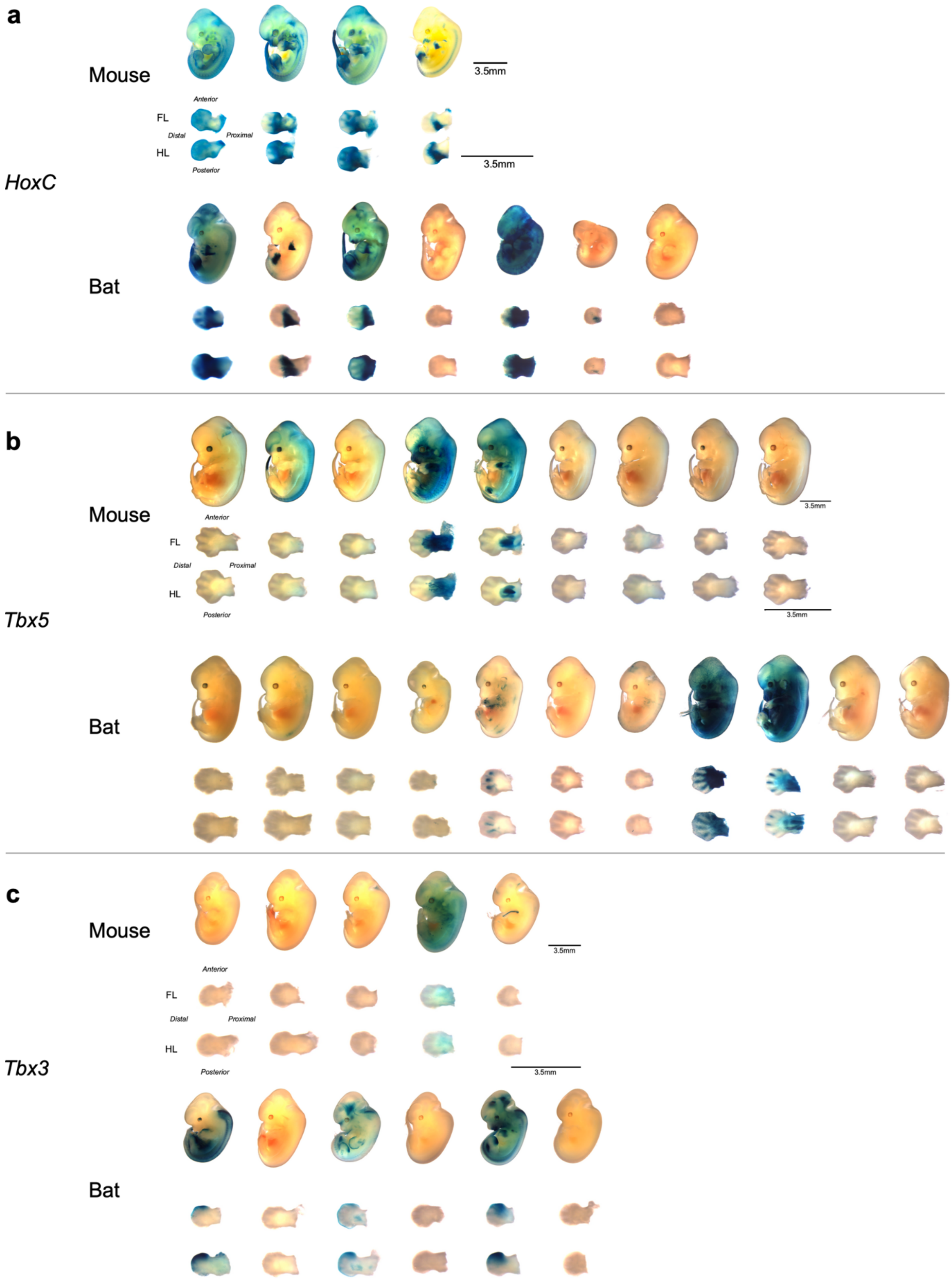


**Extended Data Fig. 2: LacZ mouse transgenic enhancer assay embryos for *HoxC*, *Tbx5*, *Tbx3***
**a-c,** Embryos and dissected forelimbs and hindlimbs for all mouse transgenic enhancer assays positive for LacZ for *HoxC* **(a)**, *Tbx5* **(b)** and *Tbx3* **(c)**. Scale bar is 3.5mm.

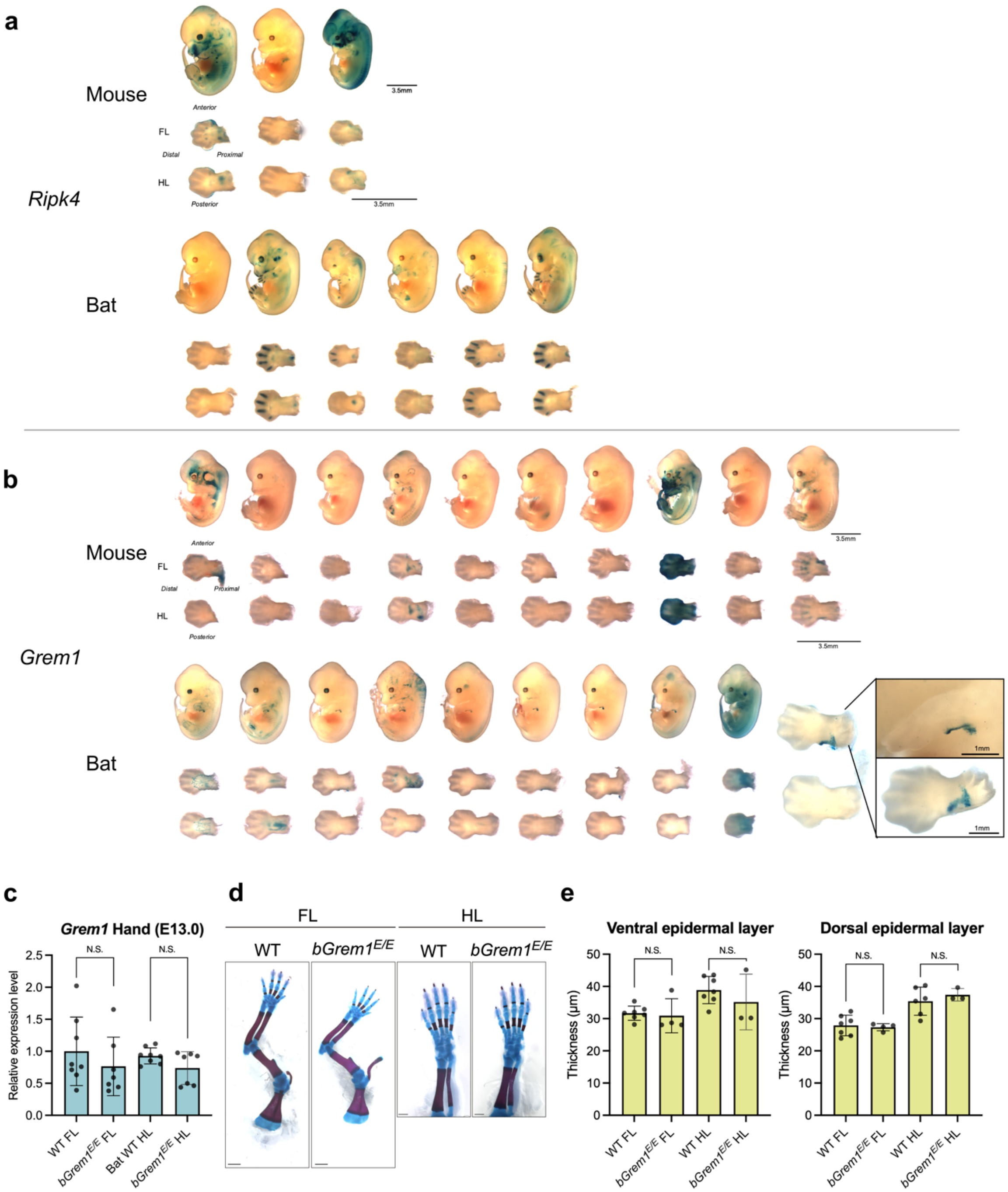


**Extended Data Fig. 3: Mouse characterization of bat *Ripk4* and *Grem1* enhancers**

**a-b,** Embryos and dissected forelimbs and hindlimbs for all mouse transgenic enhancer assays positive for LacZ for *Ripk4* **(a)** and *Grem1* **(b)**. Scale bar is 3.5mm. **(c)** Gene expression levels of *Grem1* in wild-type (WT) and bat enhancer homozygous knockin mice (*bGrem1*$^{E/E}$) in FL and HL as measured by qRT-PCR. Each value represents the ratio of *Grem1* gene expression to that of *b-Actin*, and values are mean ± standard deviation. The expression value of wild type (WT) FL was arbitrarily set at 1.0. Each dot represents one embryo. **(d)** Alcian blue and alizarin red staining for wild-type (WT) and *bGrem1*$^{E/E}$ homozygous replacement mice. Bars below denote 1cm (FL) and 1mm (HL). **(e)** Epidermal skin layer thickness in the FL and HL of WT and *bGrem1*$^{E/E}$ mice, measured in both ventral and dorsal regions at P0. Each dot represents one mouse.

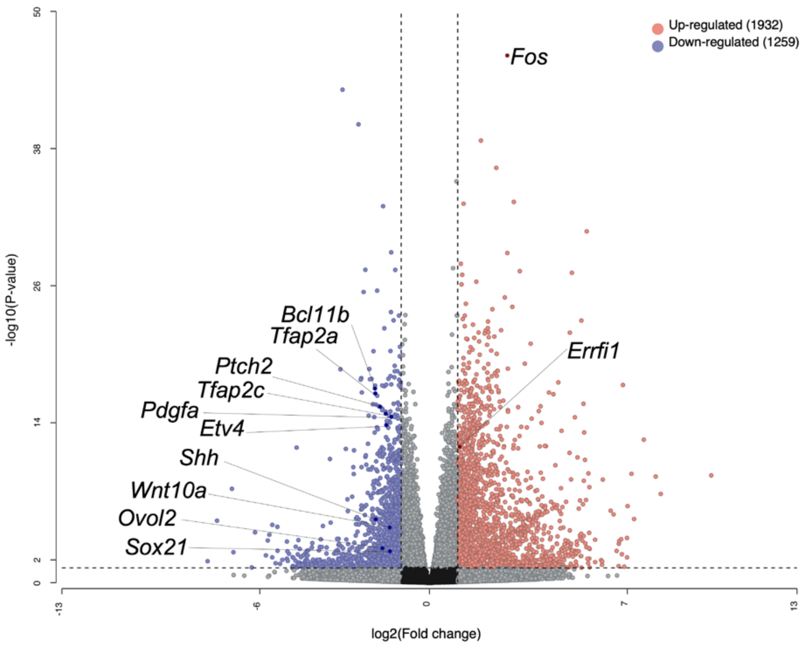


**Extended Data Fig. 4: RNA-seq of bat *Ripk4* enhancer and wild-type mice**
Volcano plots showing the global transcriptional changes from RNA-seq of *bRipk4*$^{E/E}$ versus wild-type (WT) P0 FL mouse ventral skin for the indicated groups. Each circle represents one gene. The log2 fold change in the indicated genotype is represented on the x-axis. The y-axis shows the p value. A p value of 0.05 and a fold change of 2 are indicated by lines.

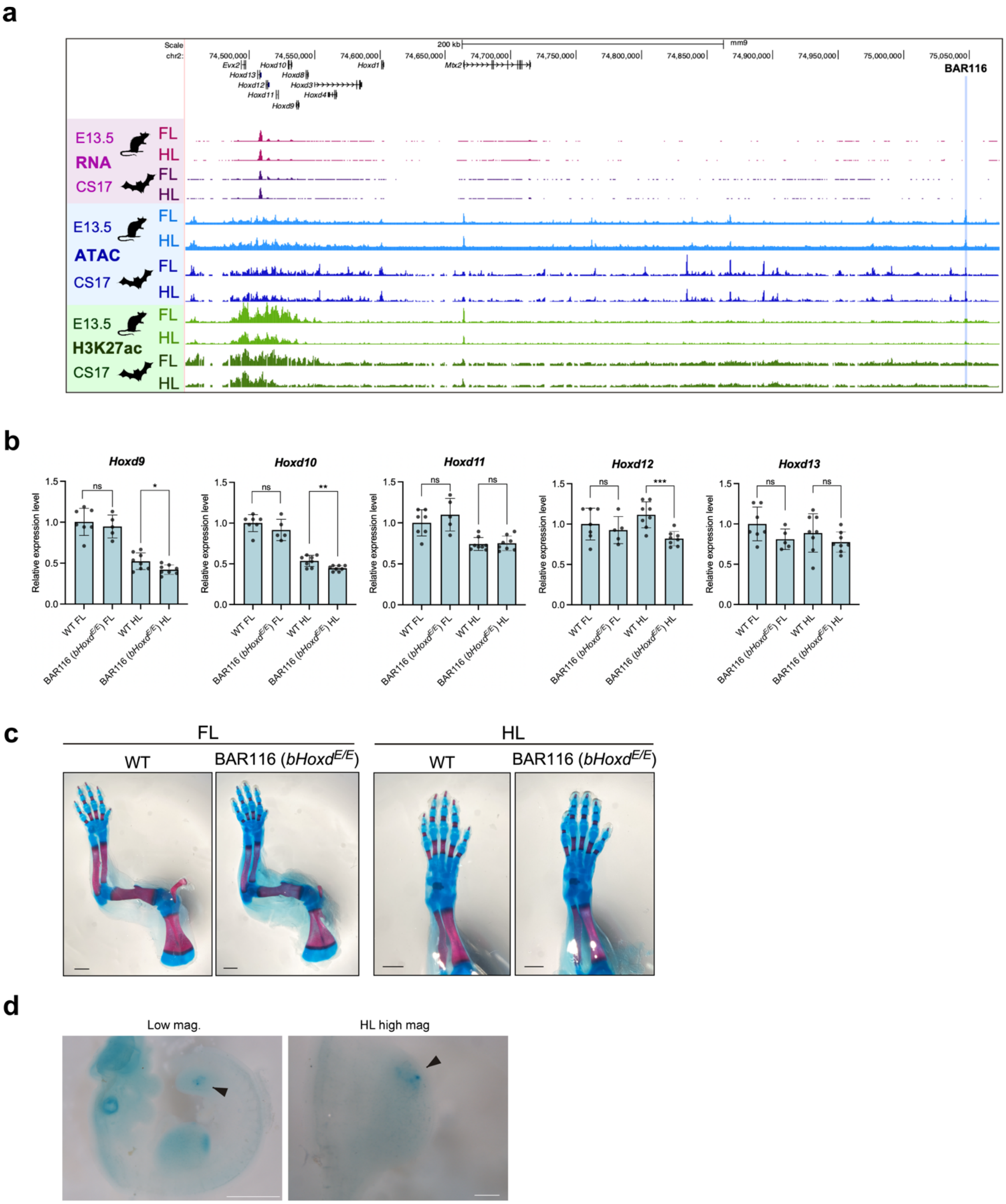


**Extended Data Fig. 5**: ***In vivo* characterization of BAR116 (*bHoxd*$^{E/E}$) and ZRS**
**a,** Genomic landscape of *Hoxd* locus. RNA-seq (pink), ATAC-seq (blue), and H3K27ac ChIP-seq (green) data from bat or mouse FL and HL at E13.5 (mouse) or CS17 (bat).
**b,** Gene expression levels of *Hoxd* genes in wild-type (WT) and bat enhancer homozygous knockin mice in FL and HL as measured by qRT-PCR. Each value represents the ratio of *Hoxd* gene expression to that

of *b-Actin*, and values are mean ± standard deviation. The expression value of wild type (WT) FL was arbitrarily set at 1.0. Each dot represents one embryo.

**c,** Alcian blue and alizarin red staining for wild-type (WT) and *bHoxd*$^{E/E}$ homozygous replacement mice. Scale bars below denote 1mm.

**d,** Whole-mount *in situ* hybridization for *Shh* of CS14 early *Miniopterus natalensis* embryo. Black bar represents 1 mm (whole embryo) and 0.2 mm (limb bud). A biological replicate of a different embryo is shown in Fig. 4F.

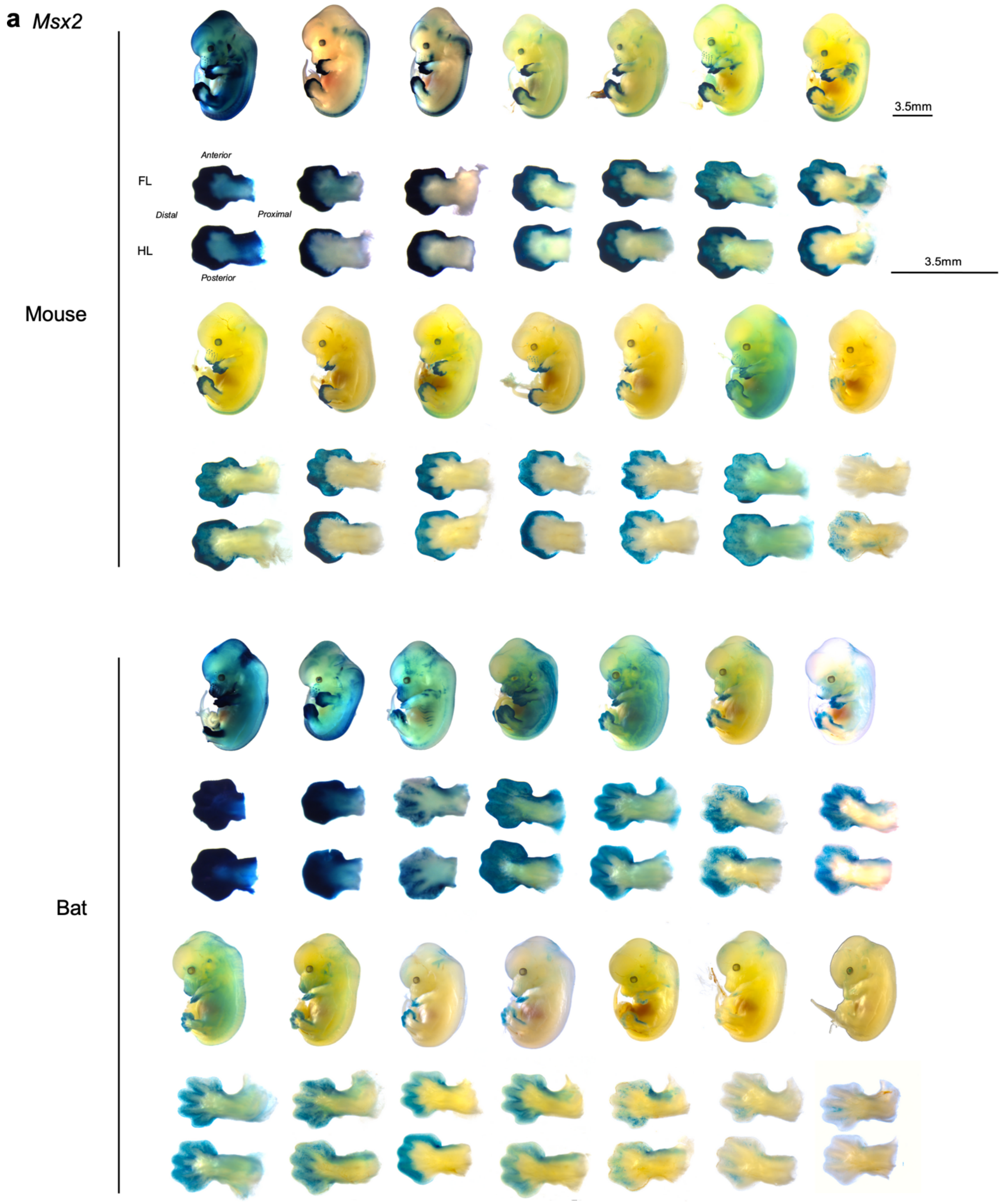


**Extended Data Fig. 6: Transgenic mouse characterization of the bat *Msx2* enhancer**
Embryos and dissected forelimbs and hindlimbs for all mouse transgenic enhancer assays positive for LacZ for *Msx2*. Scale bar is 3.5mm.

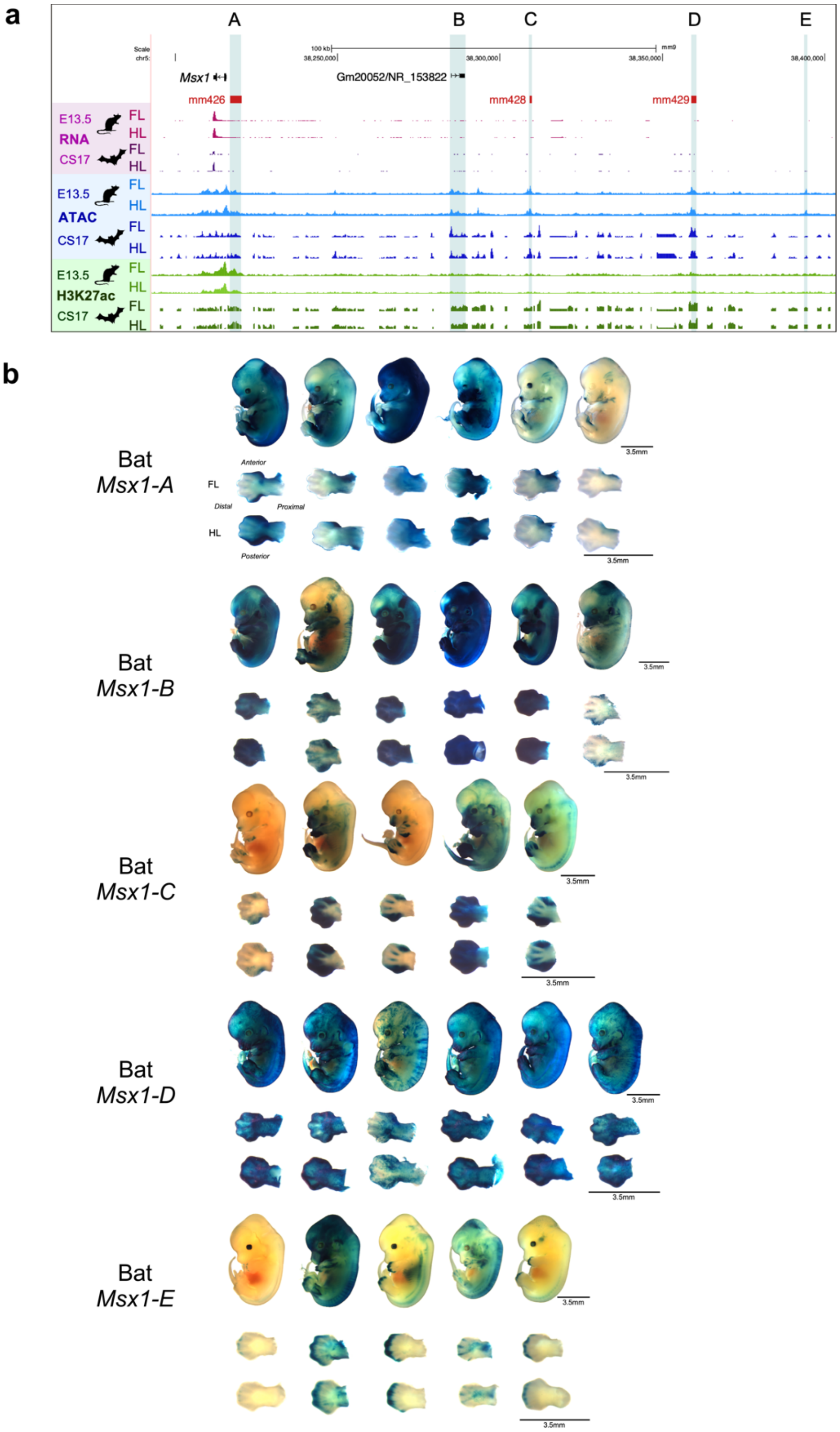


**Extended Data Fig. 7: *Msx1* locus and mouse transgenic characterization of the *Msx1* enhancer**
**a,** Genomic landscape of *Msx1* locus. RNA-seq (pink), ATAC-seq (blue), and H3K27ac ChIP-seq (green) data from bat or mouse FL and HL at E13.5 (mouse) or CS17 (bat).
**b,** Embryos and dissected forelimbs and hindlimbs for all mouse transgenic enhancer assays positive for LacZ at E13.5 for *Msx1* candidate enhancer sequences. Scale bar is 3.5mm.

a

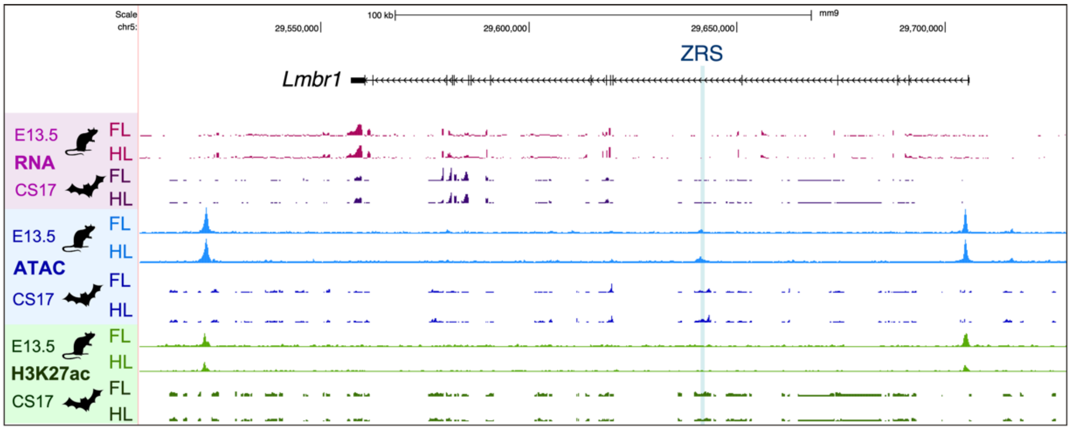


b

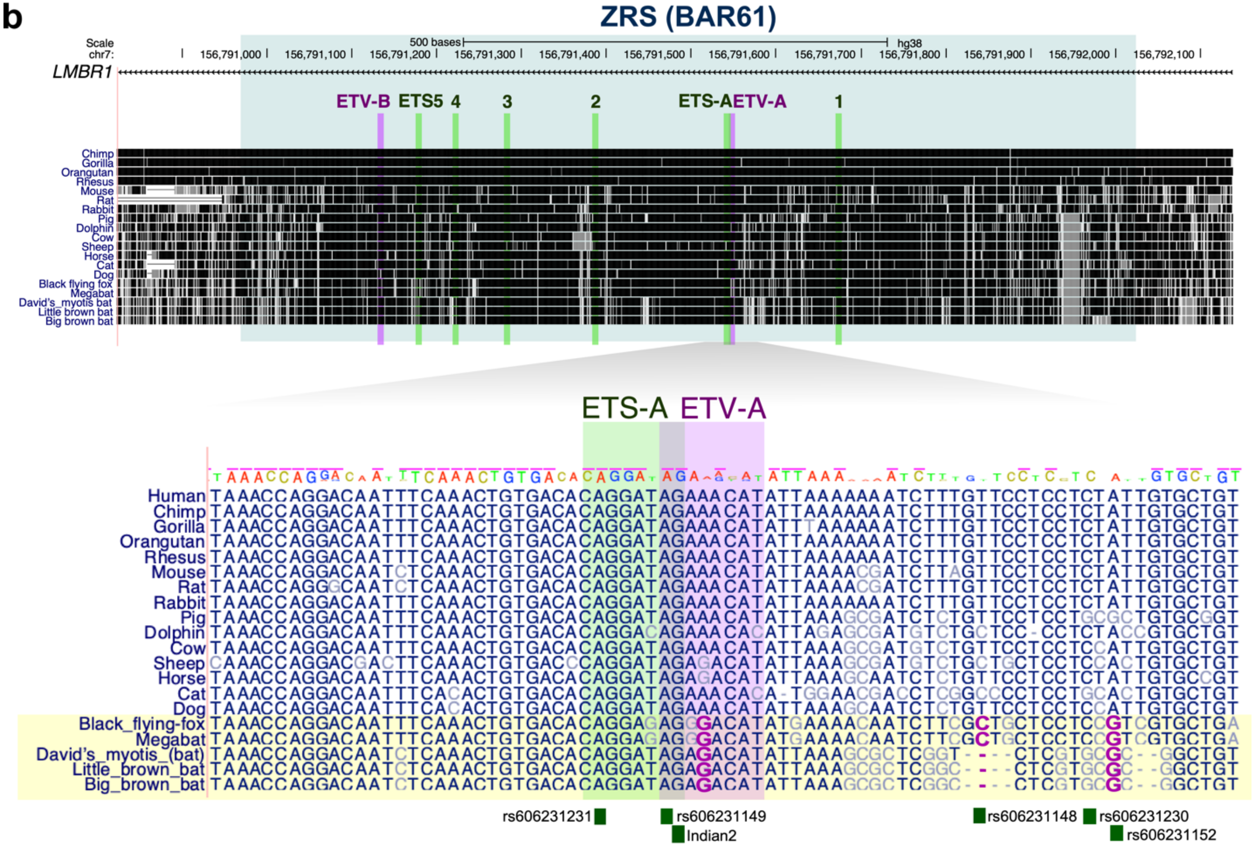


c

| Name | ClinVar allele | Condition | ClinVar Variation ID |
|---|---|---|---|
| rs606231231 | C | Polydactyly of a triphalangeal thumb | 30497 |
| rs606231149 | G | Polydactyly of a triphalangeal thumb | 4900 |
| N.A. (Indian2) | C | Preaxial polydactyly | 3061990 |
| rs606231148 | A | Polydactyly of a triphalangeal thumb | 4899 |
| rs606231230 | T | Polydactyly of a triphalangeal thumb | 30496 |
| rs606231152 | G | Polydactyly of a triphalangeal thumb | 4906 |

**Extended Data Fig. 8: ZRS sequence conservation and variants associated with triphalangeal thumb polydactyly**

**a,** Genomic landscape of *ZRS* locus. RNA-seq (pink), ATAC-seq (blue), and H3K27ac ChIP-seq (green) data from bat or mouse FL and HL at E13.5 (mouse) or CS17 (bat).

**b,** Genomic conservation of ZRS across multiple species from the UCSC Genome Browser. The bottom five species represent Chiroptera (bats). Previously reported ETS and ETV binding sites are indicated by green and purple bars, respectively. Green boxes mark the locations of human variants associated with triphalangeal thumb polydactyly.

**c,** Single nucleotide variant IDs, Condition and ClinVar(*91*) Variation IDs for relevant ZRS mutations.

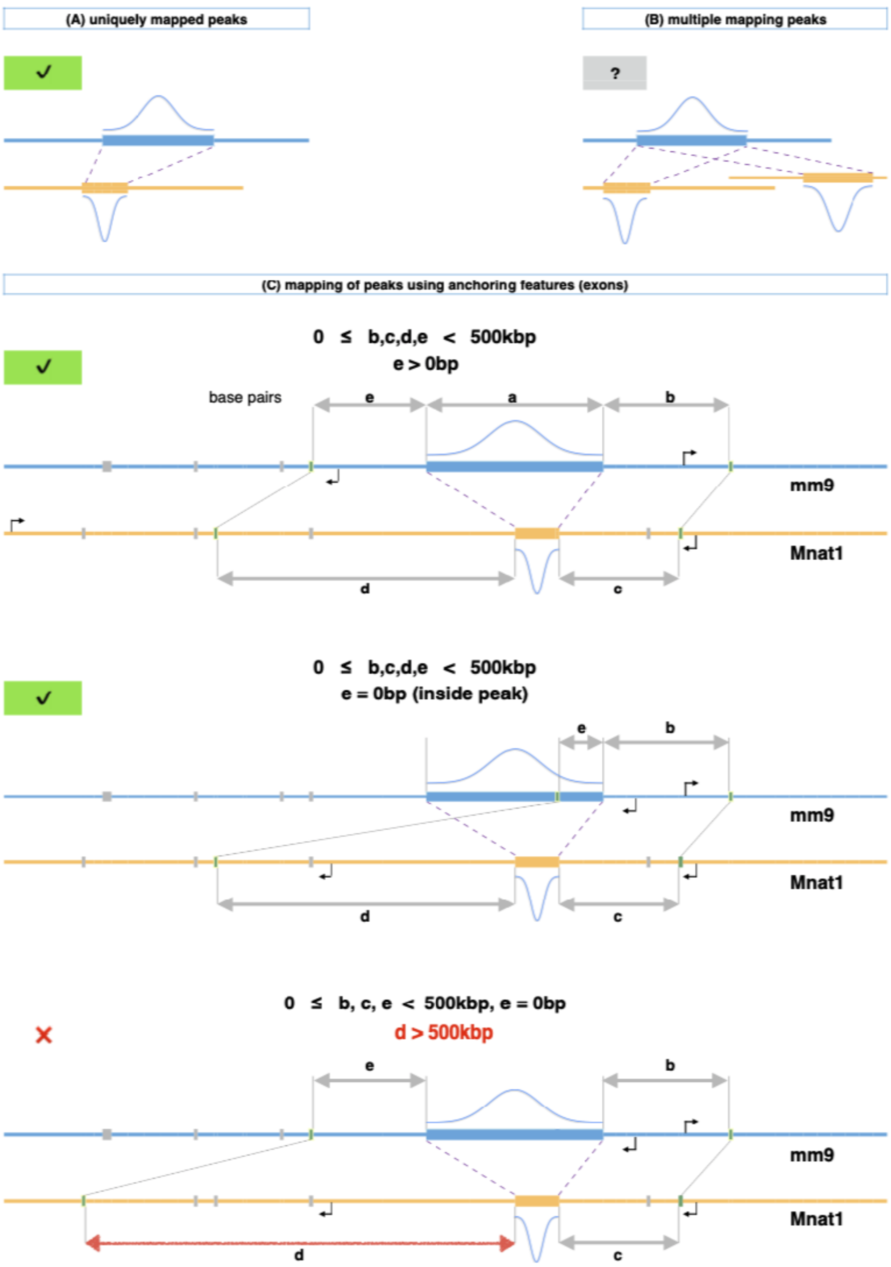


**Extended Data Fig. 9: Mapping of peaks between two species.**
Roughly one third of the peak regions are not shared between species.
**a,** Another third of the peaks uniquely map with liftOver(*92*), and no further processing is needed.
**b,** The remainder have two or more aligning candidates, and this circumstance is unresolved. For this, we utilize other synteny blocks as anchors.
**c,** Deciding a candidate peak from a multiply aligned map. The panels visualize a single mm9 ChIP peak and a single candidate peak from the Mnat assembly. Two search areas in a radius 500 kb from the peak are scanned to locate exons that map reciprocally between the species. The distances marked b-e are between the peak and its respective anchors. An anchor inside a peak is a zero-distance block. Successful mapping is marked with a green check, while the bottom panel shows an anchor on one end that is too far from the signal and the mapping fails. This method reclaims more than half of the unresolved active chromatin regions. Legend: dash lines designate aligned ranges; green marks are the anchors (aligned by the thin dotted connectors). Grey blocks indicate irrelevant (non-mapping) features.

### Supplementary Table 1. List of gRNAs

| Gene name | gRNA 1 [PAM] | gRNA 2 [PAM] |
|---|---|---|
| Bat Tbx3 | GGAGAATCTACAAGGTTAGC [GGG] | CCAAAGTCTGATCGATCAGT [TGG] |
| Bat Ripk4 | TACCTTCACAGAGGTGACGG [TGG] | CAGGACGTTGAGCTTGCTCA [TGG] |
| Bat Grem1 | CTTCTGACAAGTCAACCCCT [GGG] | GACAACTGGCCCATCTCAAT [AGG] |
| Bat ZRS | GCTGTTCTTCTGTGACGGGT [GGG] | CCTATGACAGTGCAAGTGAA [TGG] |
| Bat Msx2 | AATCACAAGCAGAGGTGTCC [AGG] | TCACCATGGCACGACCGAGA [GGG] |
| Bat BAR116 (Hoxd) | GCCACCAGCTCTGAATAGAA [AGG] | TCACACGGTAGATTTGGTAA [GGG] |
| Msx1-KO | CGGGAACAAGTAACATGATC [TGG] | CTGTTTAGAAGGTTTAGCCC [AGG] |

gRNAs are designed for FVB/NJ mouse strain

**Supplementary Table 2. List of primers**

| Mouse PCR primer | Forward 5'>3' | Reverse 5'>3' |
|---|---|---|
| Beta-Actin | GGCACCACACCTTCTACAATG | GGGGTGTTGAAGGTCTCAAAC |
| Tbx3 | TTGCAAAGGGTTTTCGAGAC | TGGAGGACTCATCCGAAGTC |
| Ripk4 | TGCGCACCTTCGACGC | GTACTCCATGACCAAGCCGAC |
| Grem1 | CCCACGGAAGTGACAGAATGA | AAGCAACGCTCCCACAGTGTA |
| Shh | CCCCAATTACAACCCCGACATC | GGCATTTAACTTGTCTTTGCACCTCTG |
| Msx2 | ATACAGGAGCCCGGCAGA | CGGTTGGTCTTGTGTTTC |
| Hoxd9 | CTGAAGGAGGAGGAGAAGCA | GTGTAGGGACAGCGCTTTTT |
| Hoxd10 | GCCAGGAGCCCACTAAAGTC | TGCTTTCCTTCTCCTGCACT |
| Hoxd11 | CCGCAGCCTCTAACTTCTACA | GCCTCGTAGAACTGATCAAAGC |
| Hoxd12 | CTCTTGCCTGCGATCTTCACT | GAATTCATTGACCAGGAATTCGTT |
| Hoxd13 | GGTGTACTGTGCCAAGGATCAG | CACATGTCCGGCTGGTTT |
